# Model Ensembling and Machine Learning Approaches to Predict the First Dose of Amoxicillin in Intensive Care

**DOI:** 10.64898/2026.02.03.703266

**Authors:** Mihály Leiwolf, Nicolas Gregoire, Sophie Magréault, Bénédicte Franck, Ombeline Krekounian, Jean-Baptiste Woillard, Vincent Aranzana-Climent

**Affiliations:** Inserm U1070 Pharmacology of Antimicrobial Agents and Antibioresistance, University of Poitiers, Poitiers, France; Inserm U1248 Pharmacology and Transplantation, University of Limoges, Limoges, France; Department of Toxicology and Pharmacokinetics, University Hospital of Poitiers, Poitiers, France; Department of Pharmacology, AP-HP, Groupe Hospitalier Paris Seine Saint-Denis, Bondy, France; Inserm UMR1137, IAME, University Paris Cité & Sorbonne University, Paris, France; 1 Univ Rennes, CHU Rennes, EHESP, Irset (Institut de recherche en santé, environnement et travail) – UMR S 1085, 35000 Rennes, France; Inserm, Centre d’Investigation Clinique 1414, Rennes, France; Department of Clinical Pharmacology, Medical University of Vienna, Vienna, Austria; Department of Pharmacology, Toxicology, and Pharmacovigilance, University Hospital of Limoges, Limoges, France

**Keywords:** therapeutic drug monitoring, population pharmacokinetics, anti-infective, precision medicine, antibiotics, mathematical modeling, quantitative pharmacology, dose

## Abstract

*A priori* model informed precision dosing (MIPD) recommends an appropriate first dose based solely on the patient’s covariates enabling faster target attainment without required concentration measurements. Population pharmacokinetic model ensembling and machine learning (ML) approaches were developed and evaluated to predict a first dose of amoxicillin in intensive care.

Following a bibliographic review, a virtual patient population was simulated based on cohorts from four published adult amoxicillin PopPK models. Model-development cohorts were reproduced, and steady-state trough concentrations were simulated using cohort-specific dosing regimens.

As reference methods, weighted model ensembling (WME) and classification tree (CT)-informed ensembling were implemented. Two novel ensembling strategies were developed: regression tree (RT)-informed ensembling, using RT to predict the log individual prediction/observation ratio, and factor analysis of mixed data (FAMD), assigning model weights based on patient similarity to original model cohorts. In parallel, four ML algorithms (support vector machine, k-nearest neighbors, random forest, and XGBoost) were trained to predict the dose achieving target concentrations based on covariates and dosing scheme. All approaches were compared with single-model PopPK dosing, standard dosing, and a nomogram, and externally validated using clinical data.

Most MIPD methods outperformed standard dosing. On simulated data, ensembling (30-42 % correct predictions) and ML (36-39 %) exceeded single-model approaches (14-32 %). RT-informed and FAMD ensembling improved performance by 6-10 % over uninformed ensembling on clinical data. In clinical patients receiving continuous infusion, ensembling further improved performance, with FAMD achieving 49 % correct predictions. ML-based ensembling eliminates the need for model selection and increase target attainment.

## Introduction

Dose individualization is essential in contexts of highly variable pharmacokinetics (PK) like critical illness.(1) Dose optimization by therapeutic drug monitoring (TDM) has demonstrated better clinical outcomes compared to standard dosing.(1) TDM increasingly relies on *a posteriori* model informed precision dosing (MIPD) which leverages population pharmacokinetic (PopPK) models to individually optimize dosing schemes based on plasmatic drug concentrations as well as the patient’s demographic and physiopathological characteristics.(2) To avoid delaying dose optimization until the first measured concentration is available, *a priori* MIPD predicts the first dose based solely on the patient’s covariates. Despite its potential clinical benefits, such as faster target attainment and fewer required concentration measurements, *a priori* MIPD remains less explored.

A single PopPK model can be used to make *a priori* predictions for every patient.(3) Model selection, however, is becoming increasingly challenging with the growing number of available models. A recent study described target attainment varying from 9 to 94 % for vancomycin models created from populations with the same sociodemographic and clinicobiological characteristics.(4) Additionally, many models are built on narrow subpopulations with specific characteristics and lack external validation. While these models may accurately describe patients similar to their development cohort, their predictive performance can be poor when extrapolated to different populations. To address this issue, Agema *et al*. developed two model ensembling methods for vancomycin and imatinib, one of which is machine learning (ML)-based.(5) At least one of these methods outperformed the best single model for each drug resulting in maximum absolute target attainments of 53.5 % for vancomycin and 78.4 % for imatinib.(5) However, the performance of the methods showed important variations from one molecule to the other. Also, using a small non-representative clinical dataset for model development can decrease their generalizability. As one of the previous methods, van Os *et al*.(6) also developed a ML-based PopPK ensembling algorithm, which selected relevant models for vancomycin using XGBoost. This approach surpassed not only the body mass index (BMI)-based empirical method, but also all single PopPK models and naïve ensembling in terms of accuracy and bias.

ML is also increasingly used not only for weight attribution, but also to directly make *a priori* predictions. For example, a combination of XGBoost, neural network, and Random Forest (RF) was trained on demographic characteristics to predict the probability of ganciclovir target area under the curve (AUC) attainment in children, enabling the iterative selection of the optimal starting dose.(7) Compared to the empirical method represented by a body surface area creatinine clearance (CRCL)-based literature equation, the ML methods resulted in higher attainment rate (25.8 % compared to 22.6 %) for ganciclovir and performed equally for valganciclovir (35.3 %).(7) In the case of vancomycin, XGBoost trained on covariates to predict AUC in neonates yielded a significantly higher target attainment rate (35.3 %) compared to the literature equation (28 %).(8) The particularity of these two applications is the development of the ML methods on a large quantity of simulated data obtained using literature PopPK models, with clinical data used solely for validation, this way increasing their generalizability.

Amoxicillin is a broad-spectrum β-lactam antibiotic, used as a first-line treatment for streptococcal and enterococcal infective endocarditis(9) and early ventilator-associated pneumonia(10). Antibiotics are widely used in intensive care (ICU), where β-lactam antibiotic underexposure increases 1.5-fold the risk of clinical failure and death.(11) However, choosing an adequate dose is challenging due to large inter-patient variability and dynamic physiopathological changes affecting drug exposure such as augmented volume of distribution and altered renal clearance.(2)

The objective of this study was to adapt the existing prediction dosing approaches and to develop new ML-based model ensembling and ML methods for amoxicillin to predict an optimal first dose in ICU to reach target concentrations based solely on covariates (*i*.*e*. before any individual concentrations are available thus *a priori*). These methods were retrospectively evaluated for their predictive accuracy in routine care data and compared to standard dose and an empiric dosing nomogram.

## Methods

### Overview

The methodology is summarized on *Figure 1*. We first simulated patient covariates and amoxicillin concentrations using publications studying the population PK of amoxicillin. Then, we split the simulations into a training and a testing dataset. Afterwards, we applied MIPD methods, some already existing and some novel (Factor Analysis of Mixed Data (FAMD) and Regression tree (RT) informed ensembling), to predict dosing regimen of amoxicillin that would attain a target based on steady-state concentrations. With those predictions, we compared their performances on the testing dataset and on a clinical dataset. Finally, we evaluated the sensitivity of the predictions to different training methods.

**Figure 1.**
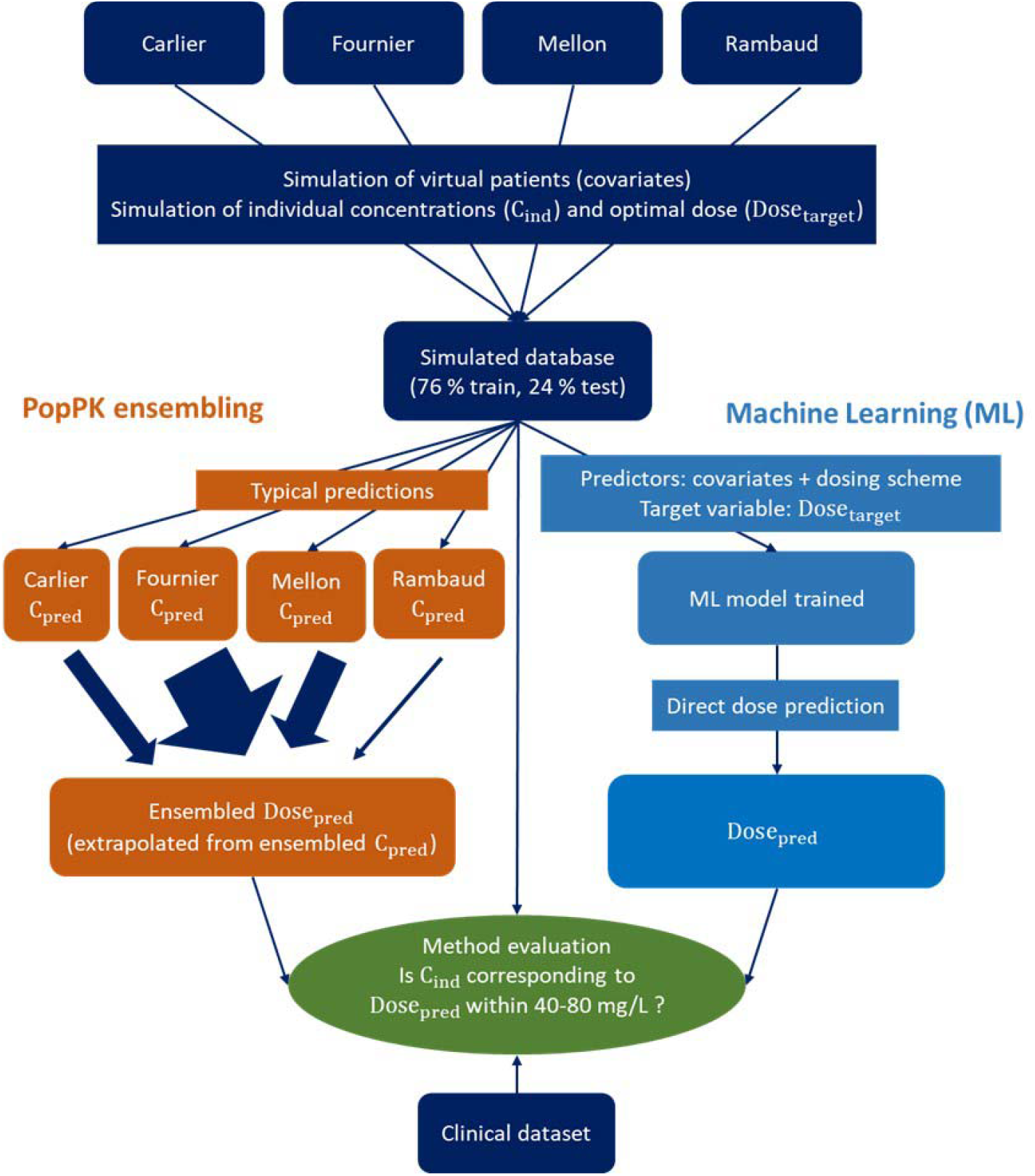
Flow chart of the methodology: First, covariates of virtual subjects are simulated. Then *true* concentrations (individual predictions – C_ind_) are simulated with each PopPK model for its own development cohort. Typical predictions (C_pred_) are made with each model for each subject, then ensembled by different PopPK ensembling methods (in orange). The ensembled prediction is converted to a predicted dose (Dose_pred_) by linear extrapolation to reach the middle of the target concentration range of 40-80 mg/L. Machine learning methods (in light blue) predict directly Dose_pred_. If this Dose_pred_ is in the interval of the *true* dose interval obtained by linear extrapolation of the true concentration to true doses of 40 (Dose_inf_) and 80 mg/L (Dose_sup_), the predicted dose is considered to be correct.

Methods are detailed in supplementary text, extensively commented R codes that enable reproduction of results based on simulated data are available as supplementary data (https://inserm-u1070-phar2.github.io/amoxicillinOpenMIPDpaper/) and dose prediction methods are available as a separate R package (*openMIPD* - https://github.com/INSERM-U1070-PHAR2/openMIPD) for use in other MIPD studies.

### PopPK models

Four PopPK models developed in adults were identified for amoxicillin. The four identified models, hereafter referred to by the name of the first author of the article, are:

- Carlier(12) developed on ICU patients
- Fournier(13) on ICU burn patients
- Mellon(14) on obese, but otherwise healthy subjects
- Rambaud(15), a non-parametric model developed on ICU patients with infective endocarditis, approximated using a parametric model by converting mean absolute weighted deviation of parameters to log-normal inter-individual variability. Rambaud was the only model developed on continuous infusion.

Model implementation was validated by the reproduction of simulation results reported in the articles (see supplementary materials).

### Database for method development

A database of 2500 virtual subjects was simulated using the four PopPK models (one cohort of 625 patients per model). For each virtual patient, covariate values were simulated, a dosing regimen was assigned based on the model cohort’s dosing schemes, and a trough concentration at steady-state simulated. Details are provided in supplementary materials.

### Simulation of concentrations and target doses

The four PopPK models were implemented in *mrgsolve* version 1.5.2. The dosing regimens corresponding to the original articles were used for the simulations with an identical number of patients for the different dosing regimens of each article (*Table S2*). The doses were adapted for renal failure patients (CRCL < 30 mL/min) when specified in the articles (Carlier and Fournier).

Steady-state trough concentrations were simulated (steady-state was considered to be reached at 3.3 times the subject-specific half-life). Individual predictions were simulated, considering inter-individual variability but not residual error, with each model for its own cohort defined by covariates and dosing regimen. Non physiological primary PK parameters (< 3 L for central volume of distribution) were resimulated with the *simeta* function. Each model used its respective formulas to estimate CRCL from serum creatinine (CREAT) and other covariates.

Target trough concentrations of 40-80 mg/L were considered for amoxicillin, according to guidelines of the French Society of Pharmacology and Therapeutics (SFPT) and the French Society of Anaesthesia and Intensive Care Medicine (SFAR)(16).

The following doses and targets were defined:

- Dose_adm_: Daily administered dose
- C_ind_: Trough concentration, considered as the ground truth and corresponding to the individual prediction (IPRED) from the PopPK model
- Dose_target_: Daily dose for which the trough concentration in the patient would be 60 mg/L (center of target interval)
- Dose_inf_: Daily dose daily dose for which the trough concentration in the patient would be 40 mg/L (lower limit of target interval)
- Dose_sup_: Daily dose daily dose for which the trough concentration in the patient would be 80 mg/L (upper limit of target interval)
- C_pred_: Predicted trough concentration for Dose_adm_
- Dose_pred_: Predicted daily dose that would yield a trough concentration of 60 mg/L

For each simulated patient, the Dose_target_ was calculated based on the administered dose (Dose_adm_) and the individual simulated trough concentrations (C_ind_) by linear extrapolation as the PK of amoxicillin was linear in all four PopPK models 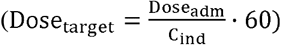. The same way, the lower and upper limits of the target dose range were calculated (Dose_inf_ to reach 40 mg/L and Dose_sup_ for 80 mg/L).

The simulated data was split into training and test sets with a ratio of 0.76:0.24. An ANOVA test was drawn to confirm that the training and test sets were statistically non different regarding covariate values (p = 0.05).

### Clinical data and external evaluation

Deidentified retrospective data of 121 observations from 74 patients (1-9 measurements/patient) collected at the surgical and neurosurgical ICUs of the University Hospital of Poitiers and at the general ICU of Avicenne University Hospital in Bobigny were used for external evaluation of the developed methods. Therapeutic drug monitoring was performed as part of routine care with samples drawn at steady state. For continuous intravenous infusion, 84 steady-state samples were drawn. For intermittent infusion, all 37 samples were collected at the University Hospital of Poitiers (24 q8h, 8 q6h and 5 q4h). For intermittent infusion, trough concentrations were measured. Each observation was treated as if it originated from an independent patient.

### Methods for *a priori* dose prediction

18 methods were compared. These methods are described in detail in the Supplementary Materials. Briefly, two of these were empirical methods: the standard dose recommended by the French scientific societies(17), and a nomogram developed for ICU patients(15).

Four were single-model methods, based on the covariate-adjusted population predictions from each of the four PopPK models considered individually, and one method was based on predictions from a PopPK meta model developed using data simulated from the four models.

Five model ensembling methods were evaluated. These approaches consist of assigning weights to the predictions from the four different PopPK models to obtain a final prediction. Among these, uninformed model ensembling method assigned equal weights to the four PopPK models; weighed model ensembling (WME) assigned weights according to the performance of the different PopPK models across covariate classes(5); one method assigned weights based on a classification tree (CT)-informed ensembling(5); another used a RT-informed ensembling; and one method relied on FAMD, assigning weights according to the proximity between an individual’s covariates and those of the patients in the four model development cohorts.

Finally, five machine-learning–based methods were evaluated, involving four different algorithms: XGBoost, random forest (RF), k-nearest neighbors (KNN), and support vector machine (SVM), as well as a Lasso-based ensembling method combining these algorithms (stacking). In addition, a final ensembling method based on a regression tree was evaluated to combine the previously described methods (Method ensembling).

### Model performance

Each method was evaluated on both the simulated test dataset and the clinical dataset. For each individual, a concentration was predicted by each of the models (C_pred_), without considering inter-individual and residual variability. For PopPK-ensembling methods, these C_pred_ were ensembled based on attributed model weights. Based on the ensembled C_pred_, the optimal dose that would result in a predicted concentration of 60 mg/L was calculated 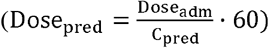.

If Dose_pred_ was in the range of Dose_inf_ and Dose_sup_, the prediction was considered correct. Thus, the performance criterion for the different methods was the proportion of subjects whose Dose_pred_ allowed for obtaining concentrations in the target interval of 40-80 mg/L.

The algorithms were not retrained on the clinical dataset; model weights were attributed and predictions were made based on the training on the simulated set.

### Sensitivity analysis

A sensitivity analysis was performed to assess the impact of different model training configurations, while testing was conducted on the same simulated and clinical datasets as previously described. The following scenarios were explored:

- Model inclusion: All combinations involving the removal of one or two models from the simulated training set were explored.
- Dosing scheme: Instead of using cohort-based dosing scheme, either a fixed intermittent dosing regimen (1 g Q8 infused over 0.5 h) or a continuous dosing regimen of 10 g was applied to the simulated training set.
- Number of subjects: One hundred datasets were generated for each of the following sample sizes: 100, 300, 600, 1200, 1900, 2500, and 3000.

## Results

### Model cohort comparison

The four simulated cohorts (see distribution details in *Table S1*, Supplementary Materials) were graphically compared to one another and to the external clinical dataset (*Figure 2*). Among them, the Mellon cohort – the only model developed on non-ICU subjects – exhibited a narrower CREAT distribution compared with both the simulated ICU cohorts and the clinical dataset, in which kidney failure and augmented renal clearance (CRCL > 130 mL/min) were more prevalent. Because the Mellon cohort was generated from obese individuals, its body weight (WT) distribution was shifted toward higher values; as a result, the simulated cohorts collectively covered the entire range of WT observed in the clinical dataset. While all subjects in the Mellon cohort were obese, the proportions of patients with a BMI above 30 kg/m^2^ were 20% in the Fournier cohort, 24% in the Rambaud cohort, and less than 1% in the Carlier cohort. In terms of age, the Mellon cohort also differed comprising fewer elderly subjects. The clinical dataset contained 55 obese subjects (45 %) and no burn patients.

**Figure 2.**
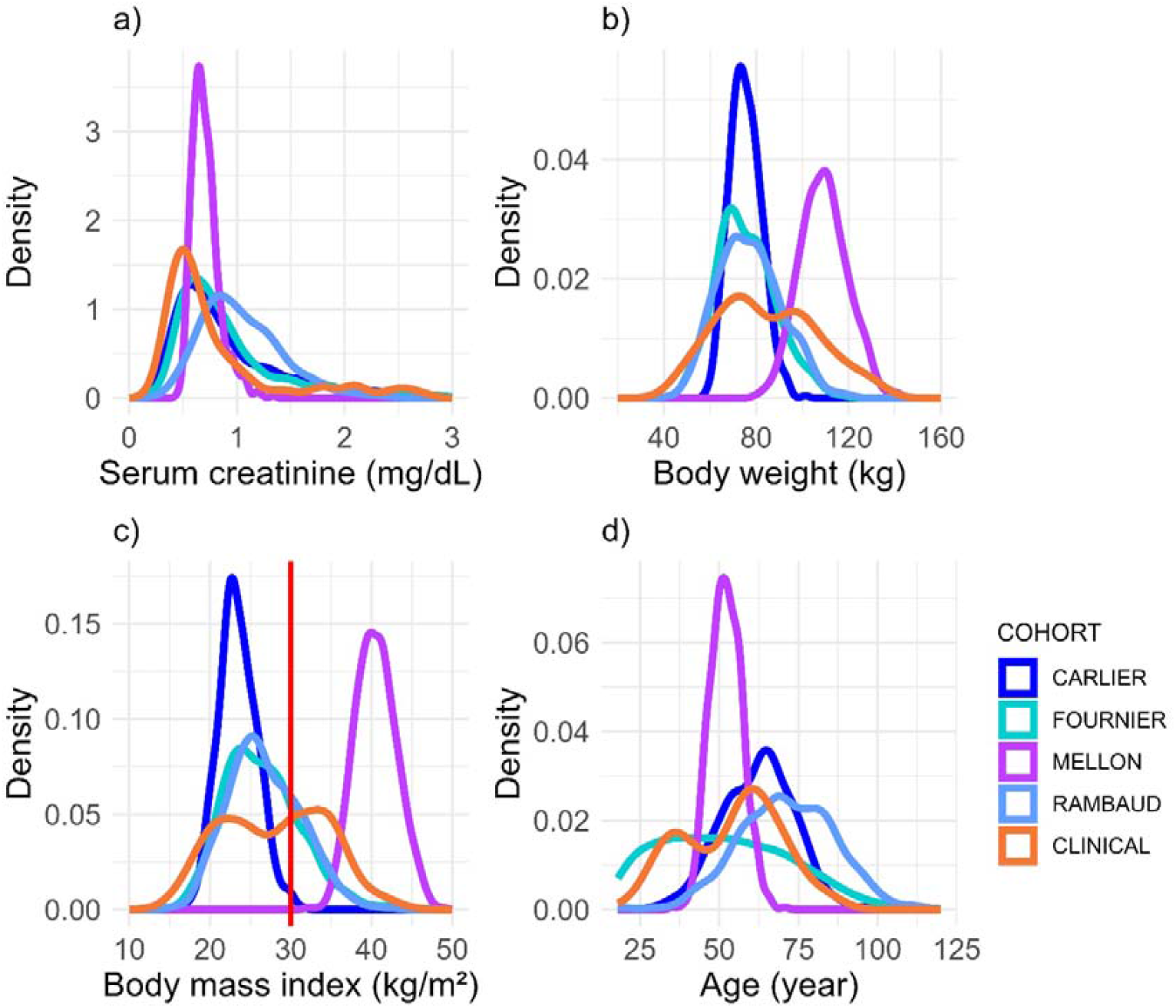
**a-d**. Distribution of serum creatinine (a), body weight (b), body mass index (c) and age (d) in the simulated training datasets corresponding to the Carlier, Fournier, Mellon, and Rambaud cohorts, as well as in the real clinical dataset. The red line at 30 kg/m^2^ on subplot c indicates the limit of obesity.

### Dose prediction on simulated test data

The PopPK meta model developed on the simulated training dataset included two compartments with linear elimination and inter-individual variability for all parameters with no covariance and a proportional error model. CREAT and WT were included as a covariate with the power model to partly explain the inter-individual variability of the clearance.

In FAMD, the seven covariates were reduced to five principal components. Among the seven covariates, ICU contributed most significantly to the model, followed by WT and OBESE (*Figure S 11-12*). For WME, the subgroup influences for the different covariates were all comparable, except for that of continuous infusion treatment, which was weaker (0.7 vs. approximately 0.85, *Figure S15*). This means that the average model performance was similar regardless of subject category, except for patients treated with continuous infusion, for whom performance was better. For RF and XGBoost, impurity-based importance analysis (18) identified CREAT, WT, age and the number of daily administrations as the most influential predictors with number of administrations on the first place for XGBoost and CREAT for RF (*Figure S16 a-b*). The method ensembling approach resulted in a decision tree only one leaf attributing XGBoost to all subjects (*Figure S16*).

When evaluated on the simulated dataset, both empirical methods – the standard dose of 200 mg/kg and the nomogram – performed markedly worse than nearly all MIPD approaches, achieving only 16 % and 22 % of correct predictions respectively (*Figure 3*, left panel). Among the four single-model approaches, Fournier demonstrated the highest *a priori* dose prediction accuracy (32 %), while Mellon reached only 14 %. In contrary to WME and CT informed ensembling, the newly developed PopPK ensembling methods (*i*.*e*. FAMD and RT informed ensembling) outperformed the best-performing single model (*i*.*e*. Fournier), meaning an absolute improvement of 4-5 %. FAMD, which does not consider model performance, outperformed the other ensembling methods. As illustrated on *Figure 3*, the single ML methods, yielded an accuracy in the range of the PopPK ensembling methods (36-39 %). Method ensembling gave an average performance of the methods it was based on (35%) rather than surpassing them.

**Figure 3.**
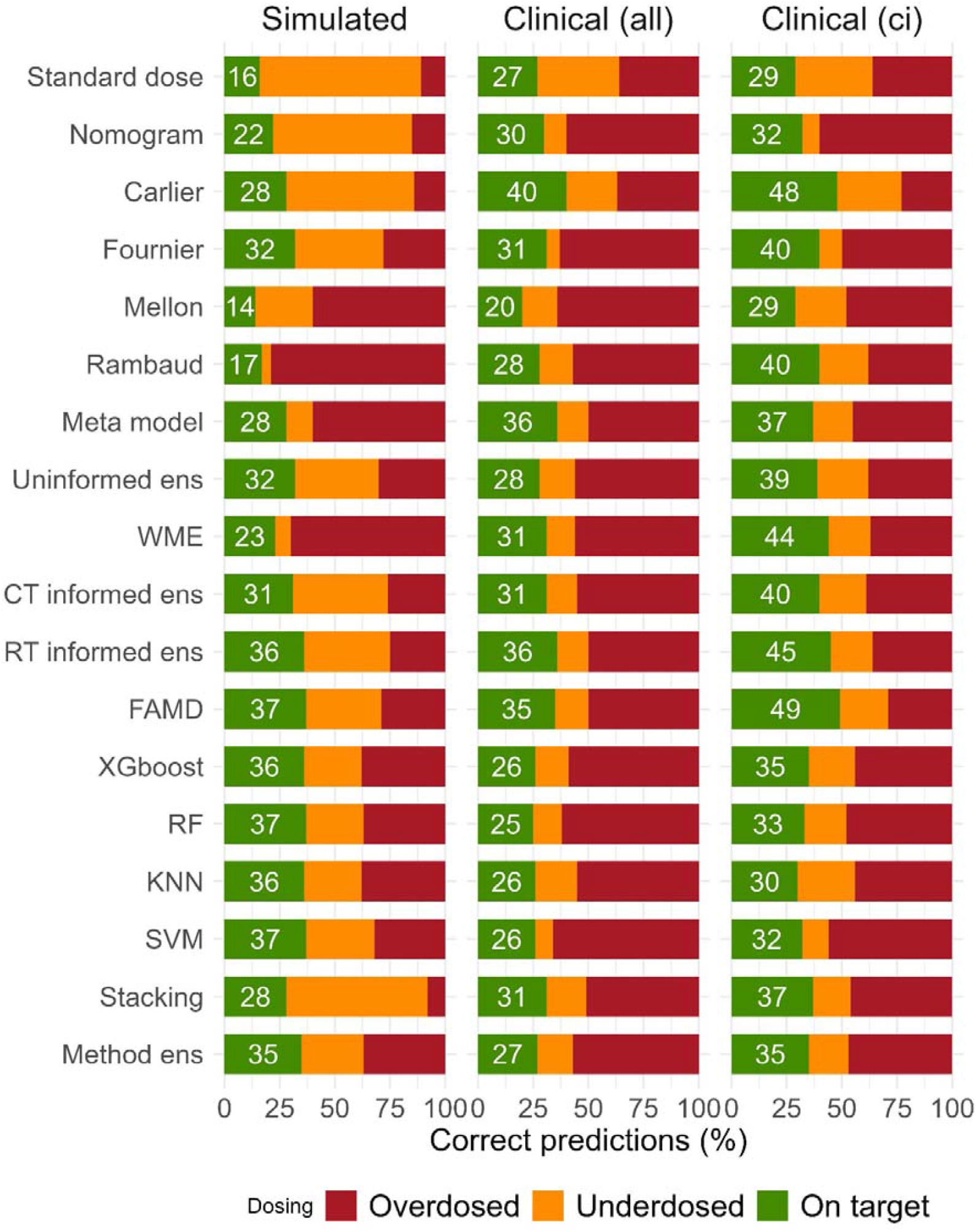
Target attainment across methods on simulated test data, clinical data, and patients in the clinical set receiving continuous infusion (ci) expressed as the percentage of patients with correctly predicted doses (green), defined as predicted doses (Dose_pred_) falling within the extrapolated dose range to achieve concentrations between 40 and 80 mg/L (Dose_inf_ and Dose_sup_ respectively). Predictions were classified as underdosed (yellow) if Dose_pred_ dose was below Dose_inf_, or overdosed (red) if it exceeded Dose_sup_.

### Clinical evaluation

The approaches trained on simulated data were externally evaluated as illustrated in the middle panel of *Figure 3*. The standard dose based on WT (*i*.*e*. 200 mg/kg), not considering infection-type (infective endocarditis or meningitis) or CRCL resulted in the highest target attainment of 27 % (*Figure S6*) performing similarly to the nomogram (30 %).

Comparing the results obtained on simulated and clinical data, a discordance can be observed. Contrarily to the simulated set, on clinical data, Carlier was the best model (*Figure 3*). The meta model developed on simulated data gained in performance and surpassed the uninformed ensembling with a performance of 36 %. FAMD and RT informed ensembling, however, showed similar predictive accuracy. The performance of ML methods decreased more markedly for clinical data, with RF showing the highest drop in performance (14 %). For the incorrectly predicted doses, overdosing was prominent compared to underdosing.

### Results for continuous infusion

An analysis of method performance stratified by infusion type (intermittent or continuous) revealed that all methods performed significantly worse on the intermittent data. When comparing the dose-normalized population predictions by the four models for clinical patients with the observed data, it appears that the models systematically underestimated concentrations following intermittent administration, whereas predictions for continuous administration were generally consistent with observed values (*Figure 4*). Given the unreliability of the intermittent infusion data due to its origin from a single hospital site, and the limited accuracy of administration and sampling time record, it is of interest to focus the external evaluation on the 84 patients who received continuous infusion, as presented on the right panel of *Figure 3*.

**Figure 4.**
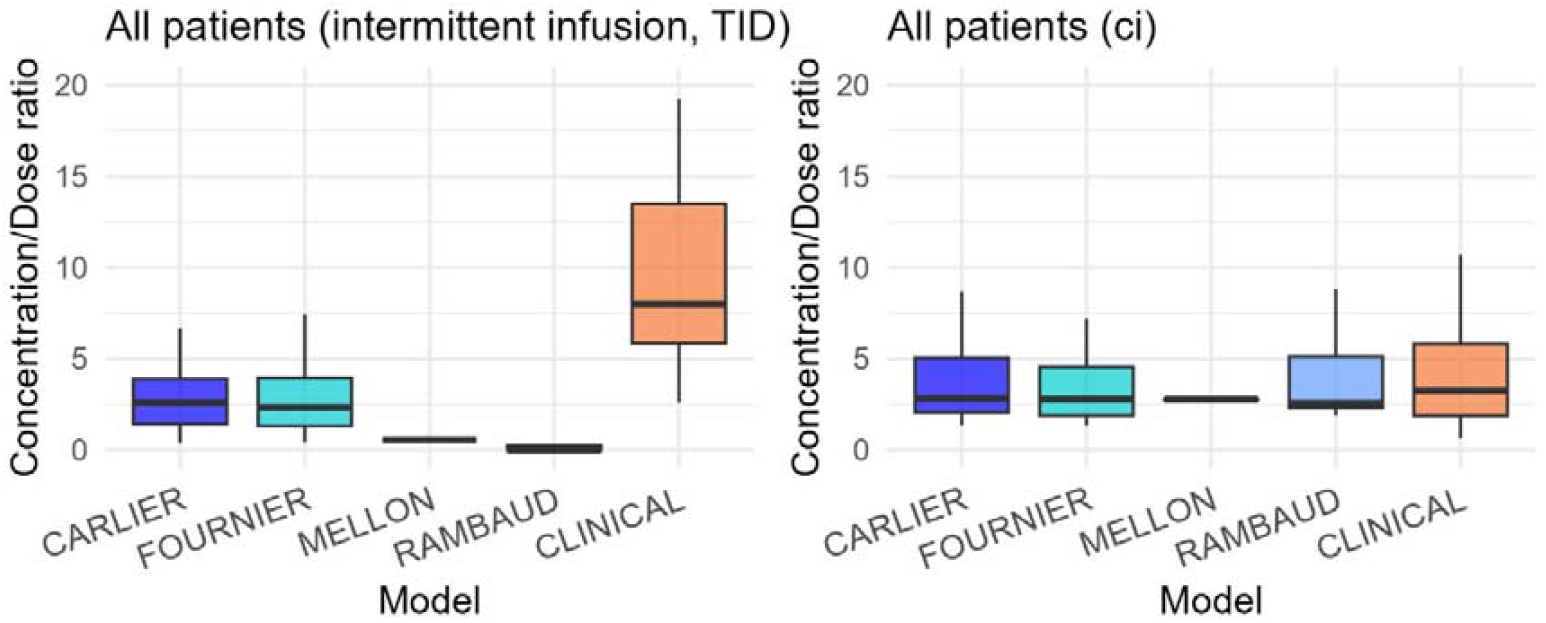
C_min_pred_ (Carlier, Fournier, Mellon, Rambaud) and measured trough concentrations (clinical) (mg/L) normalized by the dose (g) for patients in the clinical dataset who received a continuous infusion (n = 84) and for patients who received intermittent infusion three times daily (TID) (n = 24).

On data obtained after continuous administration, the results were comparable to, or even better than, that observed on the simulated test dataset (*Figure 3* right panel). Less overestimation of target doses compared with simulated data was also observed. All ensembling methods surpass the standard dose with 11-20 % of absolute improvement as well as uninformed ensembling with 1-10 %. FAMD is the best-performing method outperforming even the best single model.

As the population predictions/observed data normalized by dose illustrated on *Figure 5* shows, the variability predicted by the models, driven by incorporated covariates, and not inter-individual variability remained lower than the variability observed in the clinical dataset. It should be noted that Mellon and Fournier markedly underestimated concentrations for patients with kidney failure (CRCL < 30 mL/min). On the other hand, for obese patients, Mellon, who himself was developed from non-ICU but obese individuals, predicted concentrations comparable to other models.

**Figure 5.**
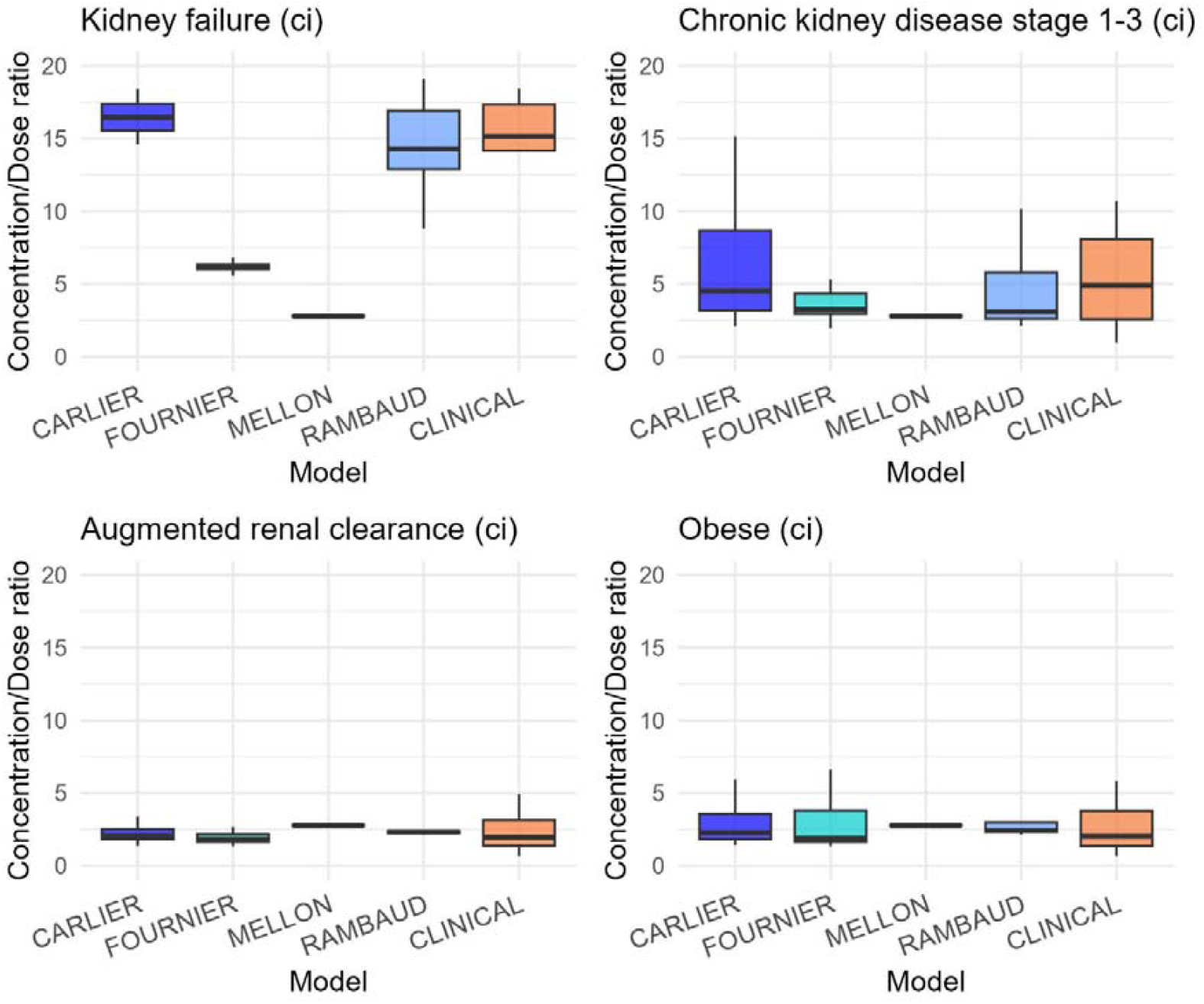
C_min_pred_ (Carlier, Fournier, Mellon, Rambaud) and measured trough concentrations (clinical) (mg/L) normalized by the dose (g) for patients who received continuous infusion (ci) stratified for obese patients, patients with kidney failure (CRCL < 30 mL/min), stage 1-3 chronic kidney disease (CRCL > 30 mL/min and < 130 mL/min), or augmented renal clearance (CRCL > 130 mL/min).

Regarding the average model weights assigned across the different patient strata on *Figure 6*, RT-informed ensembling most effectively eliminates biased and inaccurate models – specifically, Fournier and Mellon for patients with kidney failure. In contrast, WME is the least flexible, assigning similar weights across all patient groups. WME also failed to identify Carlier as the most performant method on clinical data. FAMD, which relies solely on model cohorts rather than performance, correctly eliminates Fournier and Mellon for kidney failure patients as generally assigns the most weight to the best-performing model, Carlier. However, for obese clinical patients, FAMD attributed minimal weight to Mellon as it associated obesity with not being in ICU, which shows the limitations of this method.

**Figure 6.**
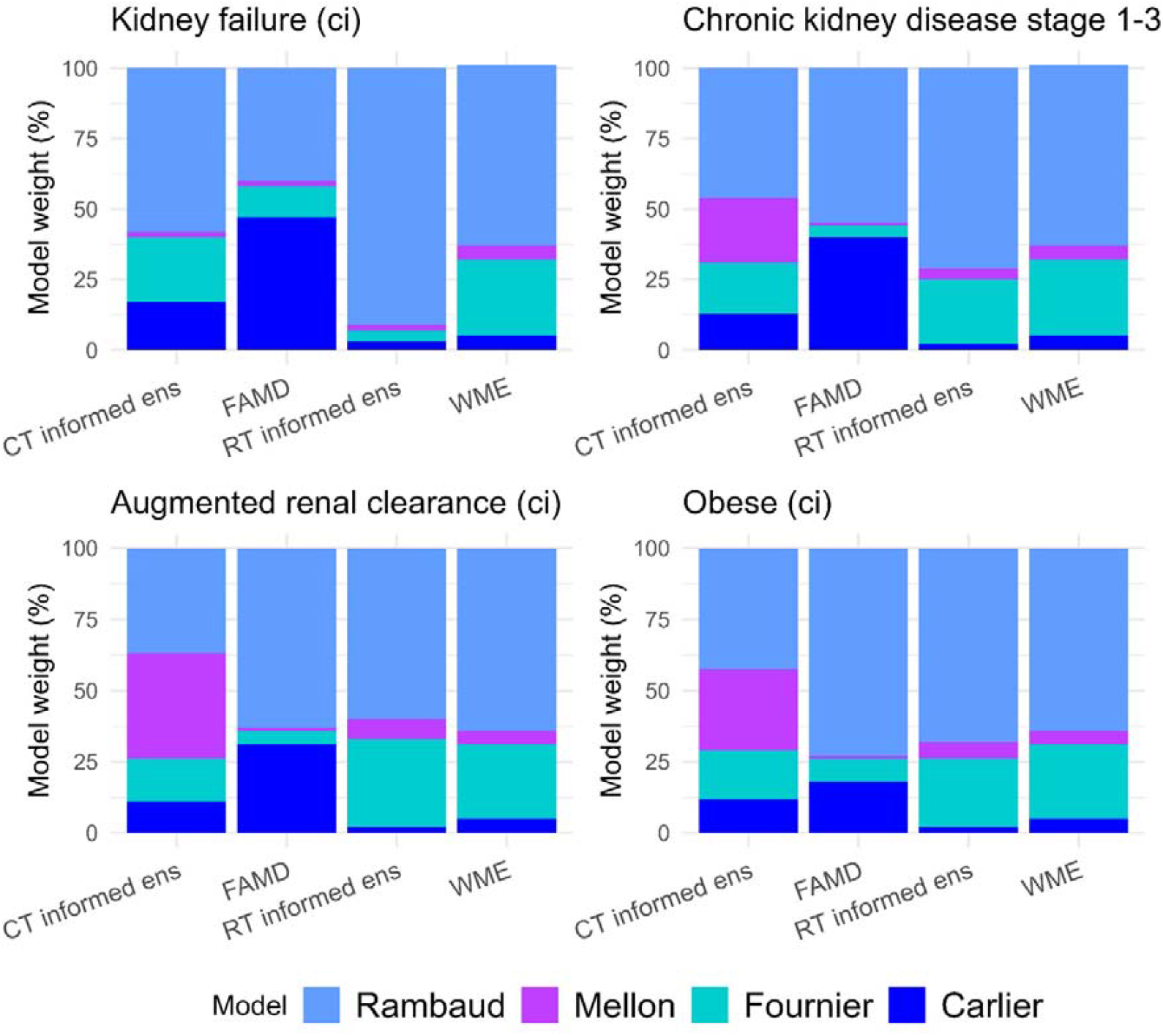
Average model weight attributed to patients receiving continuous infusion (ci) in the clinical dataset by classification tree informed ensembling (CT informed ens), factor analysis of mixed data (FAMD), regression tree informed ensembling (RT informed ens) and weighted model ensembling (WME) stratified for obese patients, patients with kidney failure (CRCL < 30 mL/min), stage 1-3 chronic kidney disease (CRCL > 30 mL/min and < 130 mL/min), and augmented renal clearance (CRCL > 130 mL/min).

### Sensitivity analysis

With respect to model inclusion, for PopPK ensembling methods, the simultaneous removal of both Carlier and Rambaud is required to noticeably deteriorate performance (*Table S3*). Removing Rambaud (i.e., the continuous infusion) results in almost exclusively overexposed predictions for the ML methods in clinical data (*Table S4*). Even though Mellon was attributed a high weight for obese patients by the PopPK ensembling methods, when it was removed in the sensitivity analysis, the proportion of obese subjects with a correct dose prediction did not decrease.

The sensitivity analysis evaluating the impact of the dosing regimens used for training showed that the best performance was achieved when the methods were trained on data simulated using the same regimens as those employed for model development (*Tables S5 and S6*). ML methods exhibited degraded performance when the dosing regimens used for training differed from those to be predicted. Conversely, training exclusively on data simulated under continuous infusion regimens improved the ability of ML to predict optimal continuous doses. PopPK ensembling methods were less affected by the choice of simulated dosing regimens than the ML approaches.

As shown on *Figures S19-20*, method performance plateaued at approximately 300 subjects in the training data, with inter-run variability (standard deviation) decreasing accordingly. A notable exception of XGBoost, which is more sensitive to overfitting on small datasets.

## Discussion

Multiple *a priori* PopPK, ML and new ML-based ensembling approaches were compared and evaluated to recommend an appropriate first dose of amoxicillin based on covariates for ICU patients. The principle of ensembling and ML approaches was benchmarked on a simulated dataset. Then, the algorithms trained on simulated data were evaluated in clinical data without retraining them.

The benefit of using MIPD was demonstrated over the best selected standard dose by ensembling methods (FAMD and RT informed ensembling). The proportion of simulated patients achieving amoxicillin trough concentrations within the 40–80 mg/L target range increased from 16 % with the standard dose to 36-37 % with PopPK ensembling. In clinical data, target attainment similarly improved from 29 % to 40-49 %. The choice of a target trough concentration range of 40-80 mg/L could be adapted to other therapeutic targets and was primarily used here as an example to enable a consistent comparison across methods.

PopPK ensembling and ML methods outperformed the single model approach in simulated data confirming the theoretical expectation that ensembling performs at least as well as, and often better than, the best individual model on which it is built. The newly developed ensembling methods (RT informed ensembling and FAMD) not only increase target attainment, but by consistently outperforming uninformed ensembling, they also eliminate the need for model selection which is complicated in clinical practice.

A discordance was observed between the simulated and clinical datasets which raises the question of extrapolability. These differences likely stemmed from the fact that the covariates in the PopPK models did not account for the variability in the observed clinical data and from the ineptitude of some models to predict dose for specific patient subgroups. For example, there were two models with biased predictions for kidney failure patients (Fournier and Mellon, *Figure 5*), whose development cohorts did not cover the whole CRCL range of the clinical data. Some methods (WME and CT informed ensembling) did not successfully eliminate these biased models by downweighing them. In addition, all models systematically underpredicted concentrations following intermittent administration. The quality of the clinical dataset – intermittent infusion originating from a single hospital and containing inconsistently documented dosing regimens and measurement times, including alternating amoxicillin and amoxicillin - clavulanic acid administrations – may have contributed to these discrepancies. When evaluated exclusively on continuous infusion, the superiority of ensembling methods over empirical approaches and uninformed model ensembling was more evidently demonstrated. These biases cannot be detected on simulated data alone, therefore external evaluation by clinical data is essential.

Training and developing algorithms on simulated data with a wide range of covariates and dosing schemes tried to address the issue of generalizability across diverse populations as well as providing a larger amount of training data. However, a key limitation of this simulation-based approach is that it relies on generating ground truth concentrations using PopPK models that may themselves be misspecified. This makes it difficult for ensembling algorithms to effectively eliminate inadequate models. For example, although in theory, all four models are bi-compartmental, Rambaud is essentially a mono-compartmental model that was developed exclusively on continuous infusion. As a result, this model exhibits parameter identifiability issues, a shorter half-life (0.211 h) compared to the other models (0.785-1.16 h) and a peripheral volume of distribution of only 0.12 L. These characteristics make it unsuitable for simulating trough concentrations under intermittent infusion. ML methods, as they lack mechanistic components, are more sensitive to training data generated with misspecified models. This was evident in the sensitivity analysis, where ML models trained on intermittent infusion could not make accurate predictions for continuous infusion, and vice versa. In contrast, PopPK ensembling methods showed greater flexibility, as the assumption of linearity in their structural models remained valid across different dosing schemes. Structural and identifiability problems of this kind were detected on simulated data where Rambaud consistently underpredicted concentrations for intermittent infusion. These issues can be resolved by simulating with each model only for the dosing schemes on which it was originally developed, and using dosing regimen as a predictor, or for ML, generate training datasets specifically simulated under the dosing regimens for which predictions are desired.

Another important question concerned model inclusion, more specifically, if a model developed on non-ICU healthy subjects (*i*.*e*. Mellon) should be used to make predictions in a critical care setting. While this model might provide valuable information for predicting optimal doses in obese ICU patients, it could also introduce bias. Our results do not clearly resolve this issue: Mellon yielded predictions comparable to those of other models, and the sensitivity analysis did not reveal any consistent improvement or deterioration in performance (on either simulated or clinical data) depending on whether it was included. Similarly, including or excluding Fournier – developed from data in burn patients, a population not represented in our clinical dataset – had no measurable impact on performance. PopPK ensembling methods that incorporate model performance appear able to appropriately weight individual model contributions and assess, for example, the informational value of the Mellon model. Nevertheless, the number of models included in training and performance evaluation is critical, as fewer models increase the risk of performance being assessed primarily on a model’s own simulated data. FAMD, in contrast, relies solely on the similarity between a patient’s covariate profile and those of the PopPK development cohorts. This makes this approach inherently dependent on the underlying model performance and inadequate to identify models with good extrapolability or bad self-predictive capacity. Notably, models with high unexplained inter-individual variability may demonstrate significant self-prediction errors (*i*.*e*. because said predictions are made using the population prediction). However, this method remains unaffected by potentially biased performance estimates derived from the training data. Interestingly, FAMD assigned little weight to Mellon for obese patients in the clinical dataset, as the ICU covariate predominated. These findings suggest that including a larger number of models in the training process may be advantageous, though this hypothesis warrants further evaluation.

The studied PopPK-based MIPD methods can easily lead to overdosing, thus, in the case of amoxicillin, increasing the risk of neurotoxicity, such as headaches or seizures. Overexposure is more important when relative bias is used (*i*.*e*. WME) as this metric favors models underpredicting the concentration. This can be improved by log-transformation of the target variable, as this way the error will represent symmetric proportional differences, not asymmetric absolute deviations. Log-transformation can be combined with using prediction/observation ratio as target variable (*i*.*e*. RT informed ensembling) instead of relative bias which further reduces dose overprediction. Log-transformation of the target variable can also improve results for ML by reducing the variance of the target.

When applied to continuous infusion (generally recommended for time-dependent antibiotics like amoxicillin), the best-performing method, FAMD, allows for a flexible, personalized weight distribution, successfully keeps the high-performing models, and demonstrates strong extrapolability correctly predicting the first dose of amoxicillin for half of real patients in unseen external validation data non representative of the training set.

Due to the lack of imputed PK information, *a priori* MIPD does not eliminate the need for at least one concentration measurement and in some cases, follow-up by *a posteriori* MIPD. Thus *a priori* MIPD is recommended as a complementary tool for the first days after ICU admission until the first available blood sample. The application of these approaches to amoxicillin is limited by the small number of available models and their similarity. The advantages of ensembling PopPK models and combining them with ML could be more easily demonstrated on a molecule with a larger number of published models with varying performance in different subgroups.

With a larger number of models, we recommend implementing all methods, as these approaches are applicable to a wide range of models. In addition, we recommend a reevaluation of these methods as an initial step, since we currently lack sufficient evidence to conclude that the methods identified as optimal in this study will systematically be the best performers.

In conclusion, ML-based PopPK model ensembling increases target attainment, enhances the robustness of predictions and overcomes the challenge of model selection. These developed methods can be readily implemented to other molecules and clinical scenarios.

## Study Highlights

1. **What is the current knowledge on the topic?** *A priori* dose selection is mainly based on empirical recommendations from scientific societies. Only few machine learning or population pharmacokinetic model ensembling methods were developed.
2. **What question did this study address?** To adapt the existing prediction dosing approaches and to develop new model ensembling and machine learning methods for amoxicillin to predict an optimal first dose in intensive care.
3. **What does this study add to our knowledge?** New machine learning-based population pharmacokinetic model ensembling methods were developed (factor analysis of mixed data and regression tree informed model ensembling), and a systematic comparison of all available *a priori* precision dosing approaches was conducted. Model informed precision dosing methods outperformed empirical recommendations.
4. **How might this change drug discovery, development, and/or therapeutics?** As the developed tools are incorporated in an R package, they are easily applicable to other molecules and clinical scenarios to perform *a priori* personalized medicine.

## Author contributions

M.L., N.G., S.M., B.F., O.K., J.B.W., and V.A.C. wrote the manuscript. M.L., N.G., B.F., J.B.W., and V.A.C. designed the research. M.L., O.K., and V.A.C. performed the research. N.G., and S.M., provided the data. M.L., N.G., J.B.W., and V.A.C. analyzed the data.

## Supplementary Information Titles

1. Supplementary Materials – Detailed methodology and additional results
2. GitHub repository https://inserm-u1070-phar2.github.io/amoxicillinOpenMIPDpaper/
3. *openMIPD* R package-https://github.com/INSERM-U1070-PHAR2/openMIPD

## Notes

**Conflict of interest** The authors declared no conflict of interest.

**Funding Information** Mihály Leiwolf was supported by an UP-SQUARED grant, (PIA4 « Excellences sous toutes ses formes » - ANR-21-EXES-0013).

Jean-Baptiste Woillard was supported by a grant from the French government, managed by the National Research Agency (ANR) under the France 2030 program, with reference number ANR-22- PESN-0017.

https://inserm-u1070-phar2.github.io/amoxicillinOpenMIPDpaper/

https://github.com/INSERM-U1070-PHAR2/openMIPD

### Competing Interest Statement

The authors have declared no competing interest.

## References

1. Wicha SG, Märtson AG, Nielsen EI, Koch BCP, Friberg LE, Alffenaar JW, et al. From Therapeutic Drug Monitoring to Model-Informed Precision Dosing for Antibiotics. Clin Pharmacol Ther. 2021/02/11 éd. avr 2021;109(4):928L41.

2. Novy E, Martinière H, Roger C. The Current Status and Future Perspectives of Beta-Lactam Therapeutic Drug Monitoring in Critically Ill Patients. Antibiot Basel Switz. 30 mars 2023;12(4):681.

3. Kantasiripitak W, Outtier A, Wicha SG, Kensert A, Wang Z, Sabino J, et al. Multi□model averaging improves the performance of model□guided infliximab dosing in patients with inflammatory bowel diseases. CPT Pharmacomet Syst Pharmacol. août 2022;11(8):1045□59.

4. Gandia P, Chaiben S, Fabre N, Concordet D. Vancomycin population pharmacokinetic models: Uncovering pharmacodynamic divergence amid clinicobiological resemblance. CPT Pharmacomet Syst Pharmacol. janv 2025;14(1):142□51.

5. Agema BC, Kocher T, Öztürk AB, Giraud EL, van Erp NP, de Winter BCM, et al. Selecting the Best Pharmacokinetic Models for a Priori Model-Informed Precision Dosing with Model Ensembling. Clin Pharmacokinet. oct 2024;63(10):1449□61.

6. Van Os W, O’Jeanson A, Troisi C, Liu C, Brooks JT, Hughes JH, et al. Machine Learning□Based Model Selection and Averaging Outperform Single□Model Approaches for a Priori Vancomycin Precision Dosing. CPT Pharmacomet Syst Pharmacol. oct 2025;14(10):1650□60.

7. Ponthier L, Autmizguine J, Franck B, Åsberg A, Ovetchkine P, Destere A, et al. Optimization of Ganciclovir and Valganciclovir Starting Dose in Children by Machine Learning. Clin Pharmacokinet. avr 2024;63(4):539□50.

8. Ponthier L, Ensuque P, Destere A, Marquet P, Labriffe M, Jacqz-Aigrain E, et al. Optimization of Vancomycin Initial Dose in Term and Preterm Neonates by Machine Learning. Pharm Res. oct 2022;39(10):2497□506.

9. Habib G, Lancellotti P, Antunes MJ, Bongiorni MG, Casalta JP, Del Zotti F, et al. 2015 ESC Guidelines for the management of infective endocarditis: The Task Force for the Management of Infective Endocarditis of the European Society of Cardiology (ESC)Endorsed by: European Association for Cardio-Thoracic Surgery (EACTS), the European Association of Nuclear Medicine (EANM). Eur Heart J. 21 nov 2015;36(44):3075□128.

10. Trouillet J. Les pneumopathies acquises sous ventilation mécanique [Internet]. SFAR; 2009 [cité 11 juin 2025]. Disponible sur: https://sfar.org/les-pneumopathies-acquises-sous-ventilation-mecanique/

11. Roberts JA, Paul SK, Akova M, Bassetti M, Waele JJ, Dimopoulos G, et al. DALI: defining antibiotic levels in intensive care unit patients: are current β-lactam antibiotic doses sufficient for critically ill patients? Clin Infect Dis Off Publ Infect Dis Soc Am. 2014;58:1072□83.

12. Carlier M, Noe M, De Waele JJ, Stove V, Verstraete AG, Lipman J, et al. Population pharmacokinetics and dosing simulations of amoxicillin/clavulanic acid in critically ill patients. J Antimicrob Chemother. 1 nov 2013;68(11):2600□8.

13. Fournier A, Goutelle S, Que YA, Eggimann P, Pantet O, Sadeghipour F, et al. Population Pharmacokinetic Study of Amoxicillin-Treated Burn Patients Hospitalized at a Swiss Tertiary-Care Center. Antimicrob Agents Chemother. sept 2018;62(9):e00505–18.

14. Mellon G, Hammas K, Burdet C, Duval X, Carette C, El-Helali N, et al. Population pharmacokinetics and dosing simulations of amoxicillin in obese adults receiving co-amoxiclav. J Antimicrob Chemother. 1 déc 2020;75(12):3611□8.

15. Rambaud A, Gaborit BJ, Deschanvres C, Le Turnier P, Lecomte R, Asseray-Madani N, et al. Development and validation of a dosing nomogram for amoxicillin in infective endocarditis. J Antimicrob Chemother. 1 oct 2020;75(10):2941□50.

16. Guilhaumou R, Benaboud S, Bennis Y, Dahyot-Fizelier C, Dailly E, Gandia P, et al. Optimization of the treatment with beta-lactam antibiotics in critically ill patients-guidelines from the French Society of Pharmacology and Therapeutics (Société Française de Pharmacologie et Thérapeutique-SFPT) and the French Society of Anaesthesia and Intensive Care Medicine (Société Française d’Anesthésie et Réanimation-SFAR). Crit Care Lond Engl. 29 mars 2019;23(1):104.

17. Lepeule R, Bru JP, Canoui E, Gauzit R, Lesprit P, Jullien V, et al. Posologie standard et forte posologie□: propositions du groupe de travail SPILF, S.P. & CA-SFM [Internet]. 2023 [cité 18 août 2025]. Disponible sur: https://www.infectiologie.com/UserFiles/File/spilf/recos/doses-spilf-sfpt-casfm-2023.pdf

18. tidyverse team. A predictive modeling case study [Internet]. tiydmodels. [cité 18 déc 2025]. Disponible sur: https://www.tidymodels.org/start/case-study/

