## Supplementary Material for "Model Ensembling and Machine Learning Approaches to Predict the First Dose of Amoxicillin in Intensive Care"

##### **Covariate simulation**

The covariates included in at least one of the PopPK models were creatinine clearance (CRCL) and body weight (WT). To characterize the relationships between covariates, correlation matrices were derived from MIMIC-IV, an intensive care (ICU) dataset comprising over 16000 patients(1). This dataset was filtered separately for each model’s development cohort by removing covariate values outside of the respective model cohorts’ ranges to ensure cohort-specific relevance. Separate correlation matrices were obtained based on sex.

For continuous variables such as WT, serum creatinine (CREAT), height (HT) or body mass index (BMI), and age, normal and log-normal distributions were tested based on the median and inter-quantile range (IQR) or minimum-maximum values reported in the original articles to select the best-fitting distribution (*Table S1* in Supplementary Materials). Using the mean and standard deviation derived from these best-fitting distributions, the correlation matrices were converted to covariances matrices to sample covariate values from a multivariate normal distribution using the log-transformed values for covariates with a selected log-normal distribution. For Carlier, standard deviation (s) for HT was obtained from the simulated log-transformed mean (m) and standard deviation of BMI and WT, and their correlation (r) based on the equations in Silverman *et al*(2):

$$\begin{aligned} m_{\mathrm{BMI}}= m_{\mathrm{WT}}-2m_{\mathrm{HT}}\#\left( 1 \right) \end{aligned}$$

$$\begin{aligned} s_{\mathrm{BMI}}^{2}= s_{\mathrm{WT}}^{2}+4s_{\mathrm{HT}}^{2}-4rs_{\mathrm{WT}}s_{\mathrm{HT}}\#\left( 2 \right) \end{aligned}$$

As CRCL was measured/estimated differently in each of the models (measured urinary clearance for Carlier, Cockcroft & Gault for Fournier, Modification of Diet in Renal Disease for Mellon, and Chronic Kidney Disease - Epidemiology Collaboration (CKD-EPI) for Rambaud), it was standardized by reconverting each CRCL estimate back to serum creatinine to ensure consistency. To account for the factors included in the CRCL formulas, age and sex were also included as covariates. As patients receiving renal replacement therapy were excluded from the original models, a minimum CRCL value of 10 mL/min was applied.

Additionally, to the four selected covariates (WT, and CREAT, sex, age), three additional categorical covariates were introduced to represent the model development cohorts:

- BURN indicating burn status, set to be true for the Fournier cohort
- OBESE indicating a BMI above 30 kg/m²
- ICU indicating models developed in ICU patients, set true for the Carlier, Fournier, and Rambaud cohorts.

***A priori* optimal dose estimation methods**

**Empirical methods**

Standard dose

Given the lack of consensus on the standard amoxicillin dosing regimen in ICU, five daily dosing schemes were evaluated:

- 100 mg/kg
- 150 mg/kg
- 200 mg/kg
- Infection-based: 200 mg/kg for patients with infective endocarditis or meningitis and 100 mg/kg for others
- CRCL-based: 100 mg/kg for patients with CRCL < 40 mL/min, 6000 mg for CRCL between 40 and 130 mL/min and 8000 mg for CRCL > 130 mL/min

These regimens were partly guided by hospital practices and partly informed by recommendations from the French Society of Pharmacology and Therapeutics (SFPT) and the French Infectious Diseases Society (SPILF)(3). The dosing regimens were evaluated on clinical data and the regimen associated with the highest target attainment was selected as a reference to the developed model ensembling algorithms.

Nomogram

A nomogram developed by Rambaud *et al*.(4) was evaluated. This method predicts the daily dose (in g) allowing to reach an amoxicillin concentration of 20 mg/L solely based on the patient’s CRCL (in mL/min) estimated using the CKD-EPI equation:

$$\begin{aligned} Dose=0.0001 \cdot CRCL^{2}+0.0613 \cdot CRCL+1.157 \#\left( 3 \right) \end{aligned}$$

The predicted dose was multiplied by three to estimate Dose_pred_ that would reach a target concentration of 60 mg/L.

Although the nomogram has been validated only for CRCL between 30 and 120 mL/min and continuous infusion, in this study, it was also evaluated in patients with CRCL outside of this range and applied to intermittent infusion.

**Single model approach**

Single models

Each model (Carlier, Fournier, Mellon or Rambaud) was used independently to predict doses for all patients in the evaluation datasets (simulated and clinical).

Meta model

As a meta-analysis of the four existing models for amoxicillin, a PopPK model was developed using Monolix 2024R1 on the training observed dataset simulated with the four original models. 1-2, and 3-compartment structural models were tested. The pharmacostatistical model was selected using the Stochastic Approximation for Model Building Algorithm. Covariates were included based on stepwise covariate modelling with respective forward and backward p values of 5 and 1 %.

**PopPK model ensembling**

Uninformed model ensembling

In uninformed model ensembling, all models were given the same weight (one quarter for each, in this case) without considering their performance or development cohort characteristics. The predictions (C_pred_, Dose_pred_) for an individual therefore corresponded to the average of the predictions of the four PopPK models.

Weighed model ensembling (WME)(5)

The training set was stratified into subgroups based on covariates (WT, CREAT, BURN, ICU, HT, SEX, AGE, and OBESE) and an additional binary variable indicating intermittent or continuous infusion (CON), with continuous variables divided into quantiles. As a result, each subject was assigned to eight different subgroups, one for each covariate.

The weighting for this method was based on the performance of the different models for the covariate subgroups and on the influence of the covariate subgroups.

The performance of each model in each subgroup was quantified as a model score. Model score was assessed using relative root mean squared error (rRMSE) and mean percentage error (MPE) calculated on trough concentrations. Model score in each subgroup was quantified by combining these metrics with an exponential function using a negative penalty value for rRMSE and MPE. The penalty values were optimized using a sensitivity analysis based on the proportion of dose predictions in the target interval.

$$\begin{aligned} rRMSE=\frac{\sqrt{\frac{\sum({C_{\mathrm{ind}}-C_{\mathrm{pred}})}^{2}}{n}}}{{m(C}_{\mathrm{ind}})} \#\left( 4 \right) \end{aligned}$$

$\begin{aligned} MPE=\frac{\sum(\frac{C_{\mathrm{ind}}-C_{\mathrm{pred}}}{C_{\mathrm{ind}}})}{n}\#\left( 5 \right) \end{aligned}$

$\begin{aligned} {Model score= e}^{\mathrm{penalty}_{\mathrm{MPE}}\cdot MPE} \cdot e^{\mathrm{penalty}_{\mathrm{rRMSE}}\cdot r\text{RMSE}}\#\left( 6 \right) \end{aligned}$

Greater influence was given to the subgroups of covariates that were most poorly predicted by the models. To identify subgroups where overall model performance was poor – indicating a need for greater attention during model selection – the proportion of $\frac{C_{\mathrm{pred}}}{C_{\mathrm{ind}}}$ratios (all models included) falling outside the 80–125 % bioequivalence range was calculated (*Figure S1*). A higher proportion of predictions outside this interval suggests that models performing well in such subgroups should be assigned greater weight in the final dose recommendation. Conversely, in subgroups where most models provided accurate predictions, model weighting was less critical. The final model weights were adjusted by multiplying model score with this subgroup-specific influence (SI) score and then normalized by their respective sum for each subject.

$$\begin{aligned} \mathrm{SI}_{CREAT-nth quantile}=\% of \frac{C_{\mathrm{pred}}}{C_{\mathrm{ind}}} \boldsymbol{\notin}\left[ 0.8;1.25 \right] for CREAT n^{\mathrm{th}}\mathrm{quantile}\left( all models \right) \#\left( 7 \right) \end{aligned}$$

$\begin{aligned} {Absolute weight}_{\mathrm{Carlier}}= \sum_{i=1}^{8} \mathrm{SI}_{i}\cdot{model score}_{i-Carlier}\#\left( 8 \right) \end{aligned}$

$$\begin{aligned} \mathrm{Weight}_{Carlier= \frac{{Absolute weight}_{\mathrm{Carlier}}}{\sum Absolute weights}}\#\left( 9 \right) \end{aligned}$$

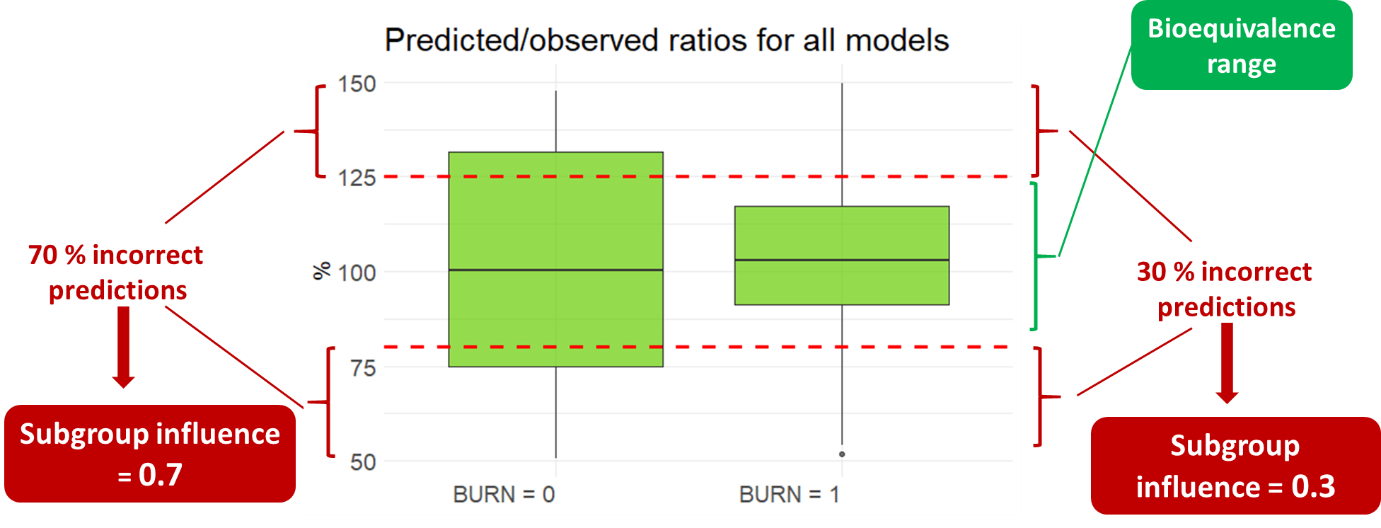

**Figure S1.** Weighed model ensembling: Subgroup influence calculation is based on the proportion of predictions, all models included, outside of the bioequivalence range of observed values. A higher proportion of incorrect predictions gives more influence to a subgroup. The figure is only illustration and does not represent results.

Decision tree (DT) informed ensembling

Classification tree (CT) informed ensembling(5)

A CT was fitted to each model using the *rpart* package in R version 4.1, with the seven covariates and CON as predictors and prediction correctness as the binary target variable (*Figure S2*). Predictions were considered correct when $\frac{C_{\mathrm{pred}}}{C_{\mathrm{ind}}}$ ratios fell within the bioequivalence range of [0.80-1.25]. The tree’s complexity parameter, which controls the size of the tree, was determined using 10-fold cross-validation, and Gini impurity was used as the splitting criterion. The minimum number of observations in a node to be split and the in a leaf (minsplit and minbucket respectively) were fixed to 4 after a sensitivity analysis. Each subject was assigned to four leaves - one for each model. In contrary to the original method developed by Agema *et al*.(8) where all models having at least 50 % of correct predictions were selected and given the same weight, in this case the proportion of correct predictions within a leaf was used to derive model weights, which were then standardized for each patient by their sum.

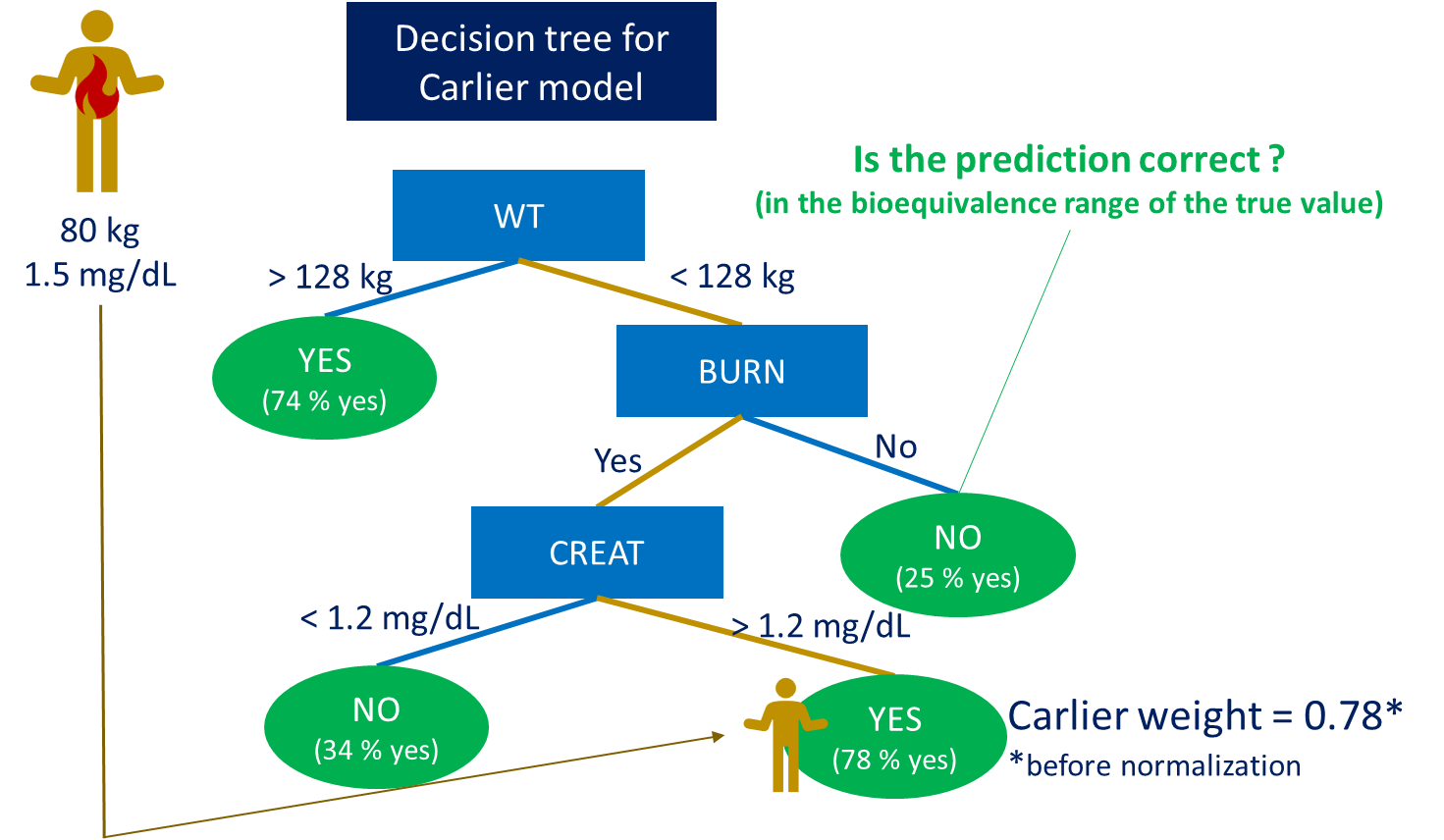

**Figure S2.** Illustration of the principle of decision tree informed ensembling. A separate tree is developed for each model. The subjects are split to nodes based on covariate values (in blue) to obtain the purest leaves (in green) where we have the highest or lowest percentage of correct (= in the bioequivalence range) predictions. If we take a specific patient as an example (in yellow), based on his covariate values, he will belong to the leaf where 78 % of patients have a correct prediction meaning that Carlier has a 78 % chance of correctly predicting the dose for him, yielding a weight of 0.78 (before normalization by the sum of the four models’ weights). The figure is only illustration and does not represent results.

Regression tree (RT) informed ensembling

This approach follows the same principle as CT informed ensembling, with the key distinction that the log-transformed prediction/observation ratio ($log(\frac{C_{\mathrm{pred}}}{C_{\mathrm{ind}}})$) was used as a continuous target variable, not prediction correctness, thereby resulting in the development of RT instead of CT. RT informed ensembling was applied using *rpart* in the *tidymodels* package with RMSE as splitting criterion. The weight assigned to each model was inversely proportional to the absolute value of the ratio shown below (example for Carlier):

$$\begin{aligned} {Model weight}_{\mathrm{Carlier}}= \frac{\left| \frac{1}{\mathrm{Ratio}_{\mathrm{Carlier}}} \right|}{\sum\left| \frac{1}{\mathrm{Ratio}} \right|}\#\left( 10 \right) \end{aligned}$$

Factor Analysis of Mixed Data (FAMD)

This hybrid approach attributes model weights based on an unsupervised ML algorithm, then applies the weighed PopPK models to make predictions. In FAMD(6) (illustrated on *Figure S3*), Principal Component Analysis (PCA) is applied to continuous covariates and multiple correspondence analysis (MCA) to categorical covariates using the *FactoMineR* package (version 2.11). The dimensionality is decreased by transforming the original data into principal components. Principal components are uncorrelated linear combinations of initial variables. They are directions in the variable space, perpendicular to each other that explain a maximum variance of the data. Principal components were added until 90 % of the variance was explained. This process simplifies the dataset while retaining most of the relevant information. Test subjects were then projected onto a latent space defined by the principal components. To determine model weights, the Mahalanobis distances(7) were calculated between each test subject and the centroids of the model cohorts within the latent space. Model weights were calculated by taking the normalized reciprocal of these distances.

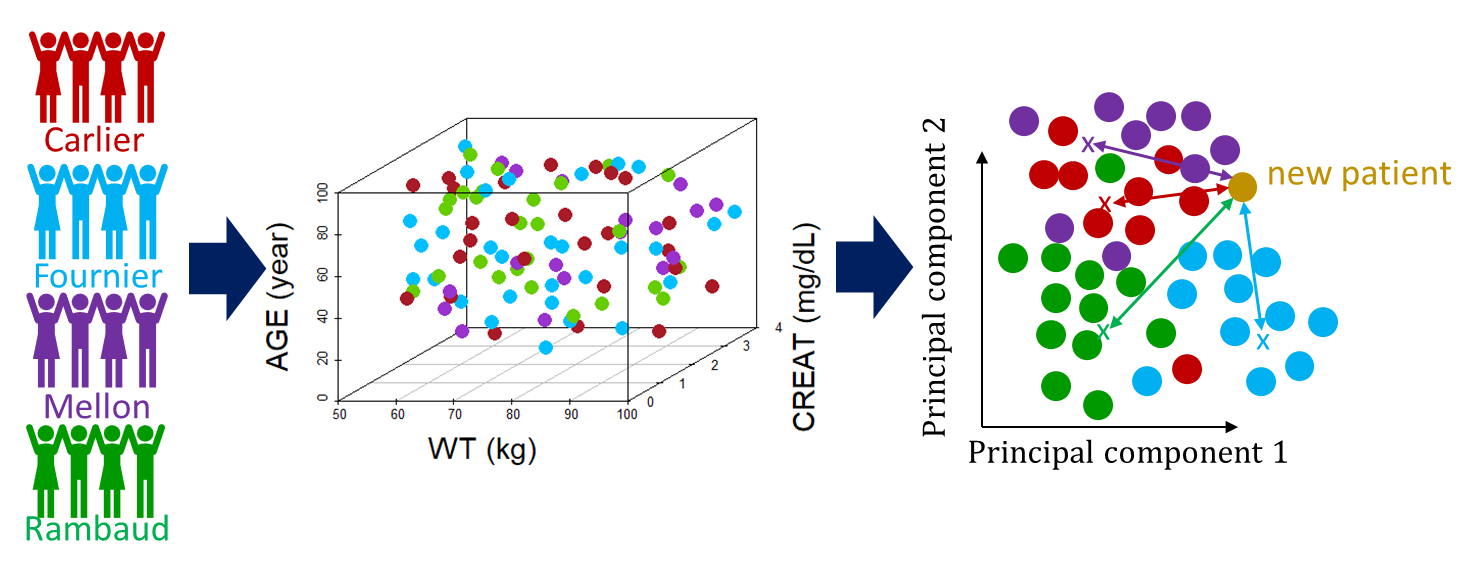

**Figure S3.** Illustration of the principle of Factor Analysis of Mixed Data (FAMD). Virtual patients are simulated based on the cohorts used to develop the original models. In this simplified example, the 3-dimensional covariate space defined by body weight (WT), serum creatinine (CREAT), and burn status (BURN) is reduced to a two-dimensional latent space through dimensionality reduction. A new patient (in yellow) is projected into this latent space, and Mahalanobis distances (indicated by arrows) are computed between the patient and the centroids (marked with x) of each model’s cohort to identify the closest matching populations. The figure is only illustration and does not represent results.

**ML methods**

Four ML methods were tested in the *tidymodels*(8) (version 1.3.0) workflow in R. As shown on *Figure S4*, these models were trained using covariates (WT, CREAT, ICU, OBESE, AGE, SEX and BURN) as predictors as well as a variable indicating the number of daily administrations to predict the dose extrapolated based on trough concentration allowing to reach 60 mg/L. For each patient, a log-transformed daily dose was predicted. Hyperparameters were tuned using 10-fold cross-validation based on mean average error and stratified by the dosing scheme.

- Random forest (RF) is based on an ensemble of decision trees. From the original training dataset, a certain number of bootstrap datasets are generated. An independent decision tree is fitted to each of the bootstrap sets with the additional randomness that at each split only a subset of the predictors is considered. The final prediction is the average of the individual trees’ predictions. RF helps reduce variance, making it suitable for unstable, unbiased data.

The hyperparameters to be defined by the user are the number of trees, the number of variables considered at each node (mtry) and the minimal number of observations in the terminal node (leaf).

- XGBoost is also a tree-based method, however, in contrary to RF, decision trees are developed sequentially to correct the prediction errors of previous trees. Model overfitting is controlled by incorporating penalized regressions, Lasso (L1) and Ridge (L2) algorithms to reduce the predictors kept in the trees. XGBoost primarily reduces bias. The hyperparameters include the learning rate that controls model parameter changes at each iteration while moving towards the minimum of the loss function, maximum tree depth and number of nodes and regularization parameters for L1 and L2.
- Support Vector Machine (SVM) classification labels data by finding a hyperplane that maximizes the distance between classes in a multi-dimensional space. SVM regression predicts a continuous target by finding a function that allows small errors within a defined margin while penalizing larger deviations from the true values. SVM regression can capture non-linear relationships and is less sensible to outliers compared to linear regression.

Common hyperparameters are the tolerance for errors (ε) and a regularization parameter (cost).

- k nearest neighbors (KNN) makes predictions by identifying the *k* closest points in the feature space and estimating the target based on their values, typically using a distance metric such as Euclidean distance.

Hyperparameters are *k* and the metric used to calculate distance between the data.

Additionally, a ML ensembling method was evaluated:

- Stacking from *tidymodels*: Predictions from the four ML models were combined by fitting a Lasso model to the data stack, with weights assigned based on the Lasso coefficients.
-
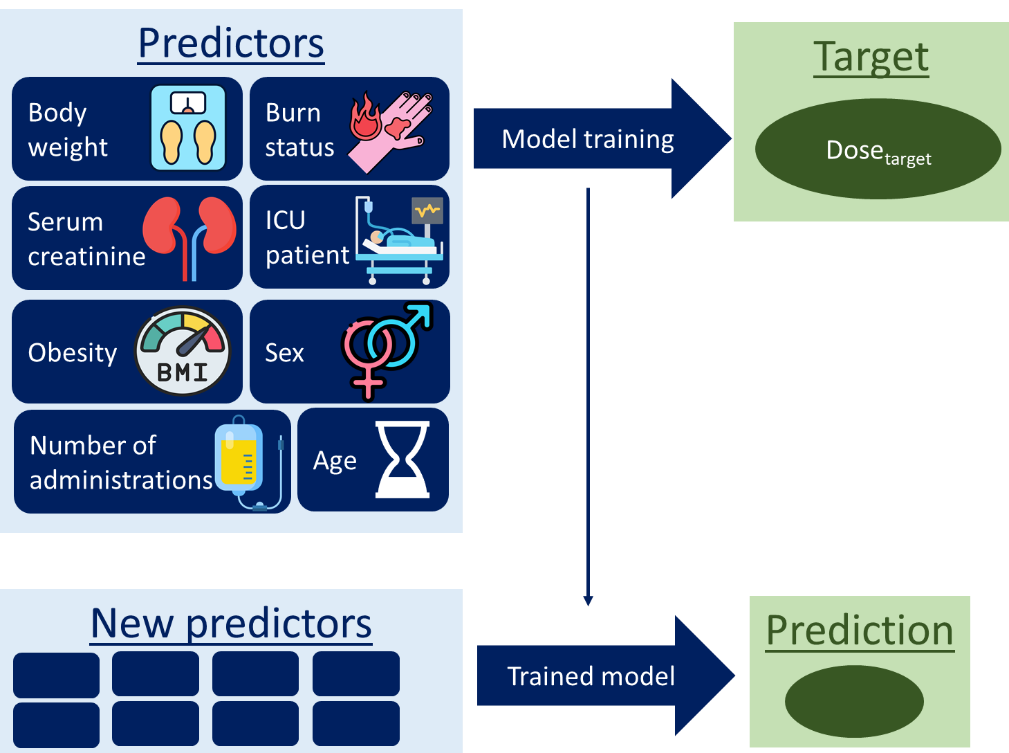

- **Figure S4.** Illustration of the principle of ML model training. Covariates and number of administrations (blue cases) are used as predictors and the target dose to reach 60 mg/L obtained by linear extrapolation from trough concentrations is used as target variable (green circle).

Method ensembling

To identify the most suitable ensembling (WME, CT informed ensembling, RT informed ensembling, and FAMD) or ML (SVM, KNN, RF, XGBoost) method for each patient and to evaluate covariate-specific performance patterns, a decision tree was developed. For each patient, the best method was selected, which yielded the closest (smallest difference) Dose_pred_ to Dose_target_. A decision tree was fitted with the seven covariates as predictors and the best method as a categorical variable comprising 8 possible classes (1 for each ensembling or ML method).

#### Simulations

**Table S1.** The simulated cohorts’ continuous covariates’ (Creatinine clearance (CRCL) and body weight (WT), height (HT), serum creatinine (CREAT), and body mass index (BMI)) selected distribution and simulated median values with the original reported median as a reference. For Mellon, CRCL was given as mL/min/1.73 m², which was converted to mL/min after the simulation of body surface area based on WT and body mass index with a correlation of 0.892.

| **Model cohort** | **Covariate** | **Unit** | **Selected distribution** | **True median** | **Simulated median** |
| --- | --- | --- | --- | --- | --- |
| Carlier | CRCL | mL/min | Normal | 102 | 115 |
| Carlier | WT | kg | Log-normal | 75.0 | 74.7 |
| Carlier | BMI | kg/m² | Log-normal | 24.0 | 23.2 |
| Carlier | AGE | year | Normal | 62.0 | 63.0 |
| Fournier | CRCL | mL/min | Normal | 128 | 117 |
| Fournier | WT | kg | Log-normal | 72.4 | 74.4 |
| Fournier | AGE | year | Normal | 50.1 | 50.6 |
| Mellon | CRCL | mL/min | Log-normal |  | 93.9 |
| Mellon | WT | kg | Log-normal | 109 | 109 |
| Mellon | AGE | year | Normal | 51.1 | 52.2 |
| Rambaud | CREAT | μmol/L | Log-normal | 87.0 | 86.7 |
| Rambaud | WT | kg | Normal | 76.0 | 76.5 |
| Rambaud | AGE | year | Normal | 72.0 | 70.8 |
| Rambaud | HT | cm | Normal | 170 | 169 |

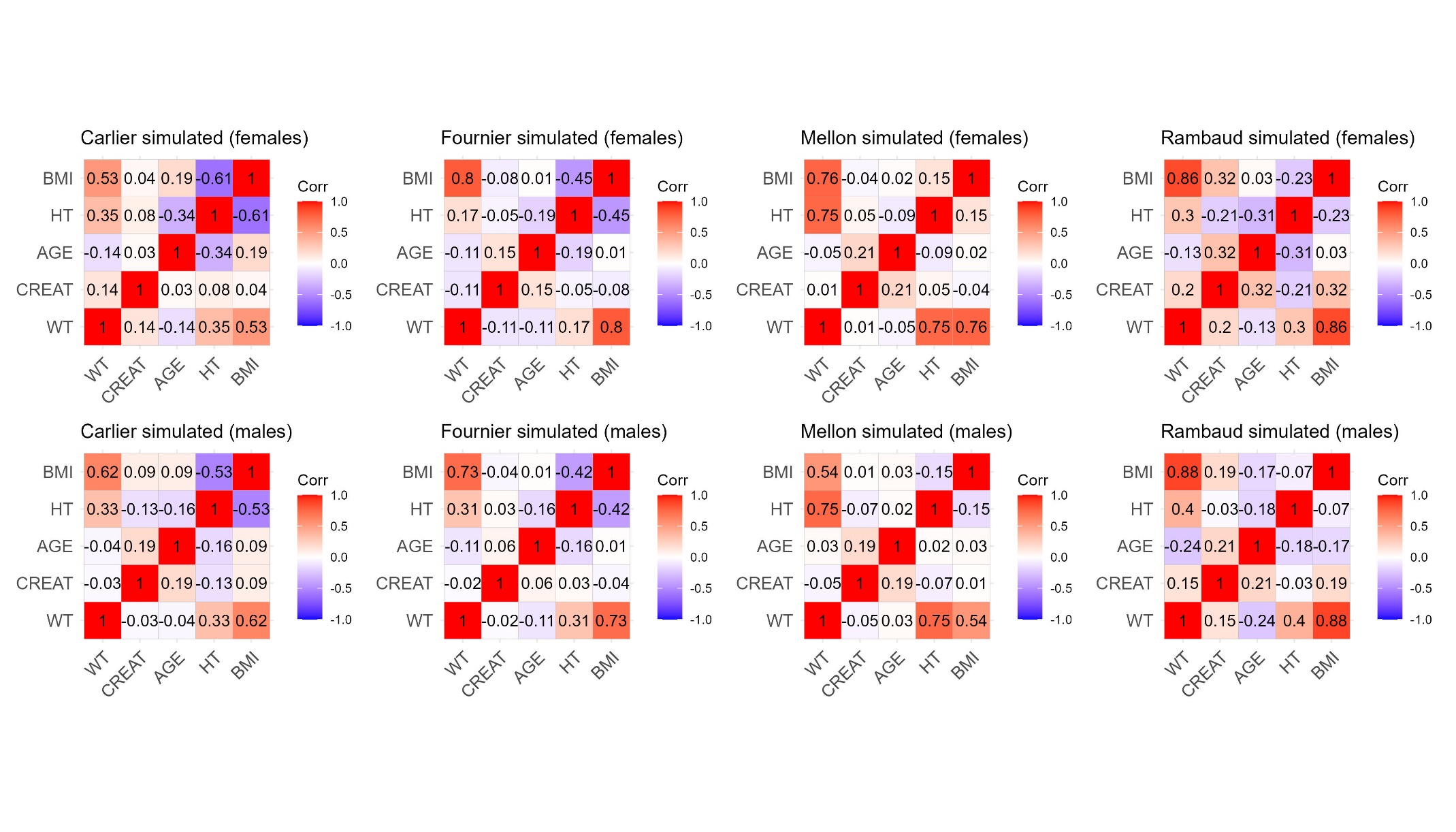

**Figure S5.** Pearson correlation plots for the simulated covariates for each model cohort, stratified based on sex.

**Table S2.** Simulated dosing schemes based on dosing schemes found in the original articles distributed based on model cohorts and CRCL values. For Rambaud, continuous infusion was administered, while for the other models, intermittent/extended infusion. For Mellon, CRCL was originally in mL/min/1.73 m².

| **Model** | **Infusion duration (h)** | **Interdose interval (h)** | **Dose (mg)** | **CRCL (mL/min)** |
| --- | --- | --- | --- | --- |
| Carlier | 0.5 | 6 | 1000 | ≥ 30 |
| Carlier | 0.5 | 8 | 1000 | < 30 |
| Fournier | 2 | 6 | 1000 | ≥ 30 |
| Fournier | 1 | 6 | 1500 | ≥ 30 |
| Fournier | 2 | 6 | 2000 | ≥ 30 |
| Fournier | 1 | 8 | 2000 | ≥ 30 |
| Fournier | 2 | 8 | 500 | < 30 |
| Mellon | 0.5 | 6 | 1000 |  |
| Rambaud | 24 | 24 | 12000 |  |
| Rambaud | 24 | 24 | 14000 |  |

#### Standard dose

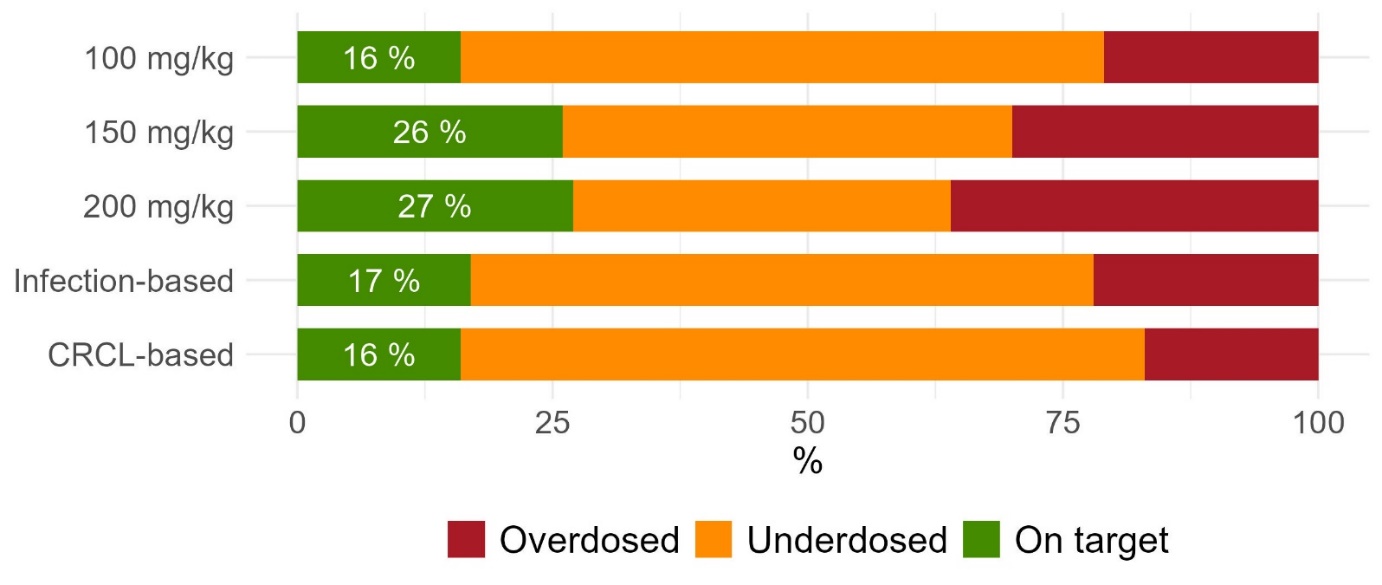

**Figure S6.** Target attainment across standard dosing schemes on clinical data, expressed as the percentage of patients with correctly predicted doses (green), defined as predicted doses (Dose_pred_) falling within the extrapolated dose range to achieve concentrations between 40 and 80 mg/L (Dose_inf_ and Dose_sup_ respectively). Predictions were classified as underdosed (yellow) if Dose_pred_ dose was below Dose_inf_, or overdosed (red) if it exceeded Dose_sup_. Infection-based = 200 mg/kg for patients with infective endocarditis or meningitis and 100 mg/kg for others. CRCL-based = 100 mg/kg for patients with CRCL < 40 mL/min, 6000 mg for CRCL between 40 and 130 mL/min and 8000 mg for CRCL > 130 mL/min. The doses are daily doses.

#### Classification tree-informed ensembling

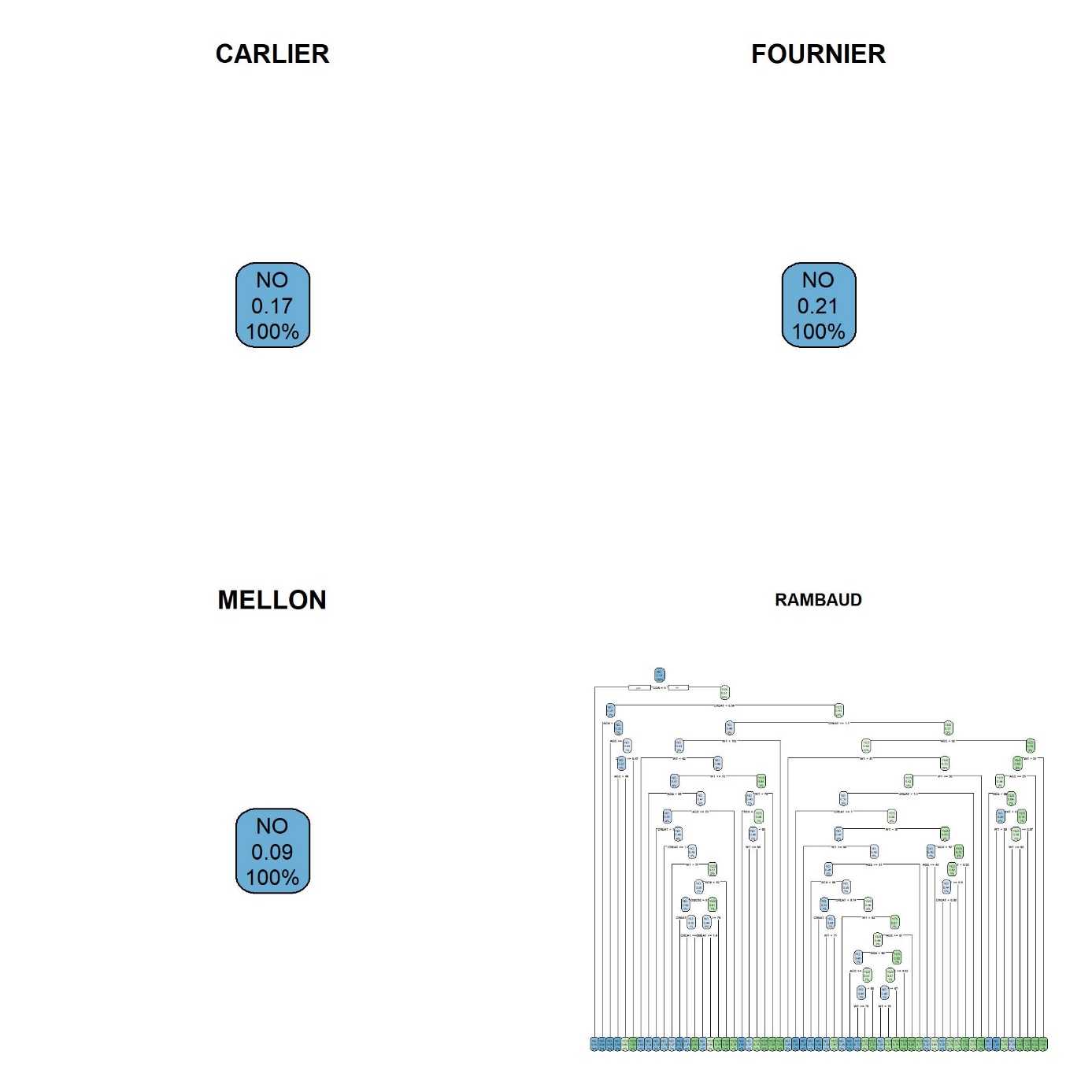

**Figure S7.** Classification trees for Carlier, Fournier, Mellon, and Rambaud. On the visualized nodes, the first line indicates the majority class in that node meaning if the majority of C_pred_ are in the bioequivalence range of C_ind_ or not (YES/NO), the second line the proportion of correct predictions (= proportion of YES), and the third line the percentage of subjects in that particular node.

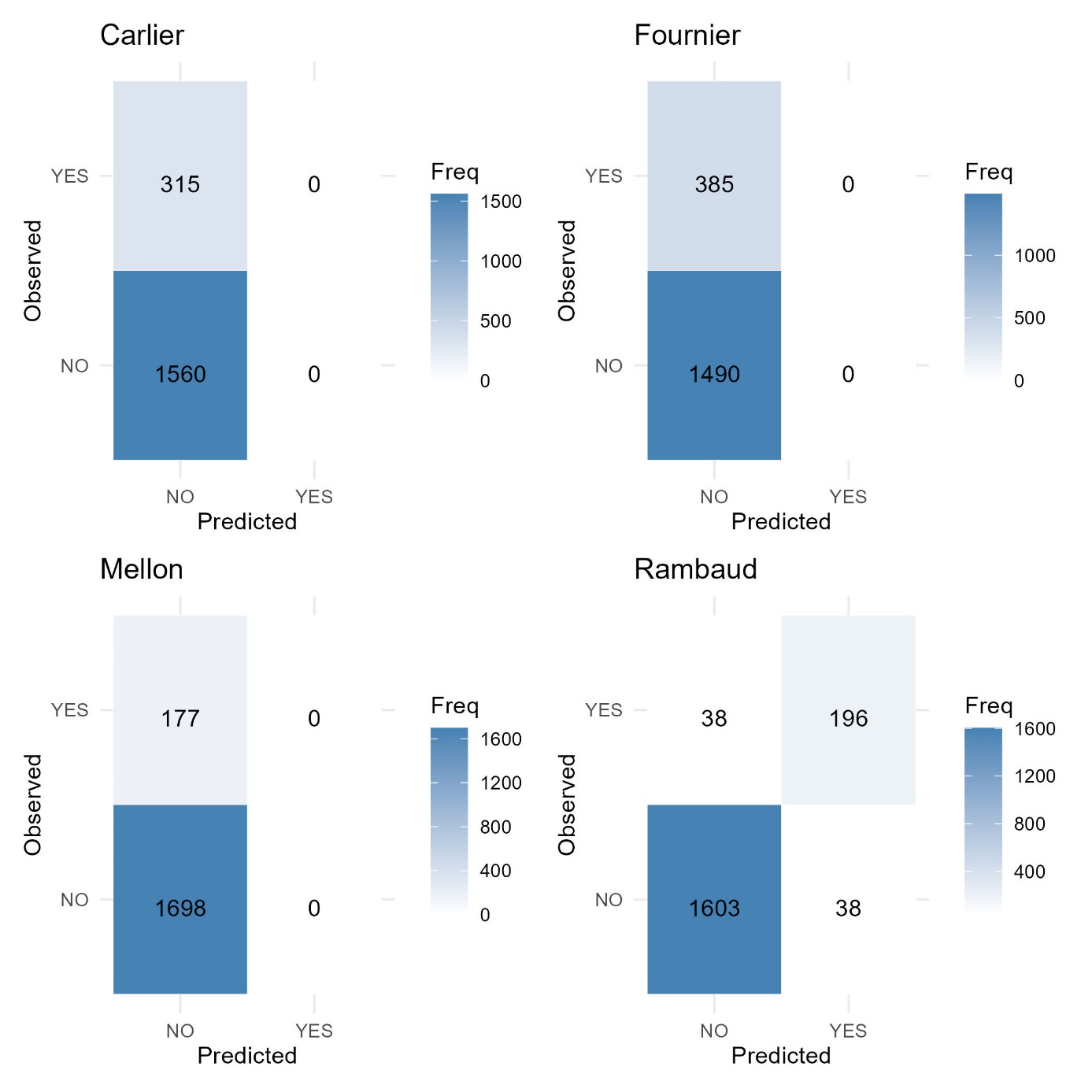

**Figure S8.** Confusion matrices for the classification trees (shown in *Figure S7*) for Carlier, Fournier, Mellon, and Rambaud.

#### Regression tree-informed ensembling

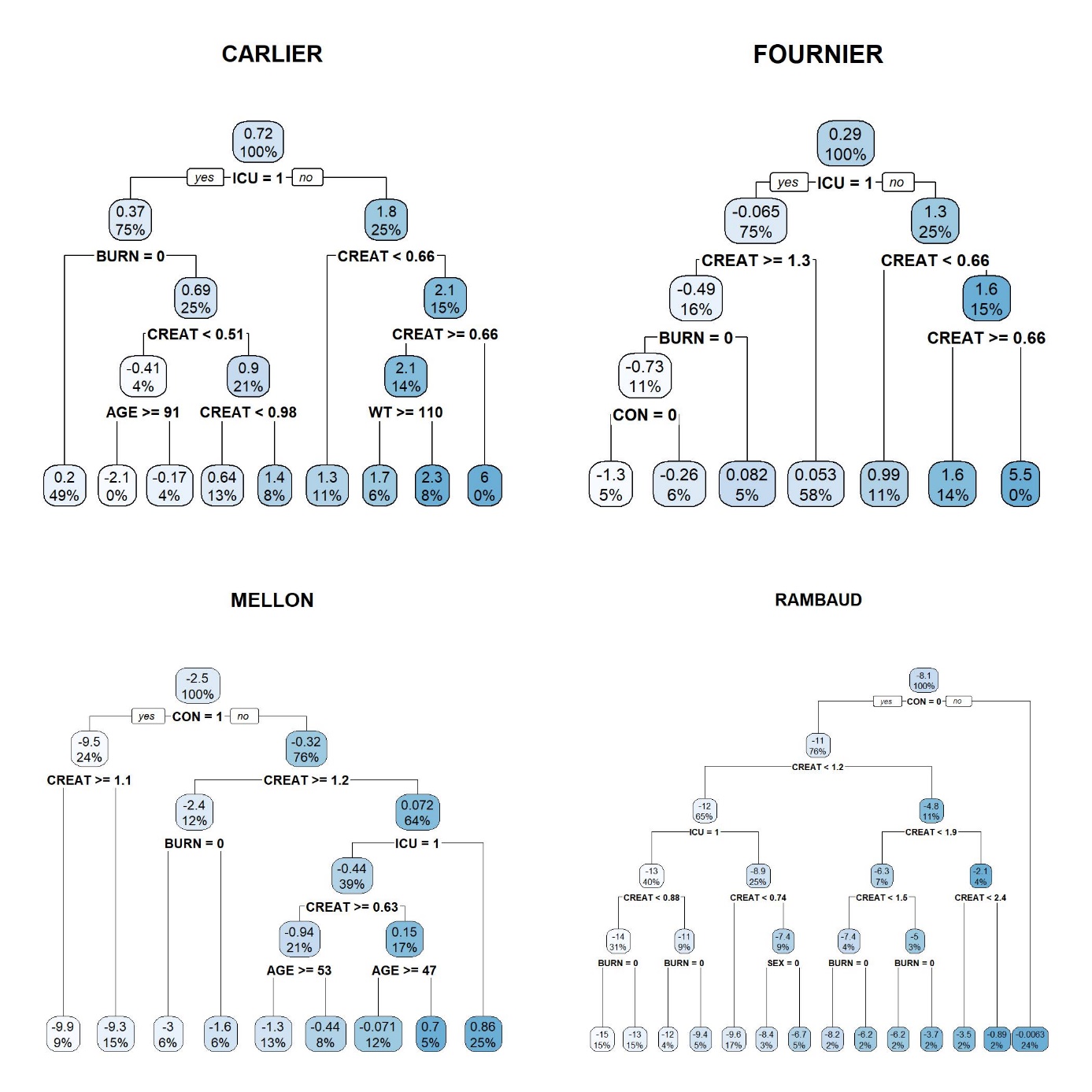

**Figure S9.** Regression trees for Carlier, Fournier, Mellon, and Rambaud. On the visualized nodes, the first line indicates average log-transformed C_pred_ / C_ind_ ratio and the second line, percentage of subjects in that particular node.

#### Factor analysis of mixed data (FAMD) egression-tree informed ensembling (RT inf ens)

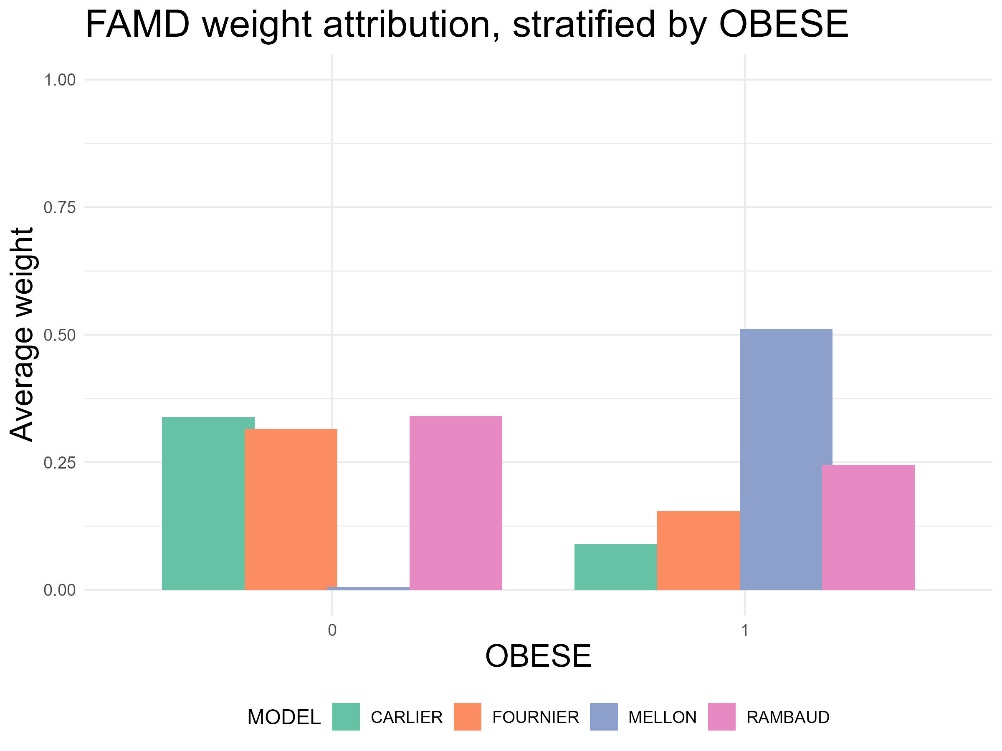

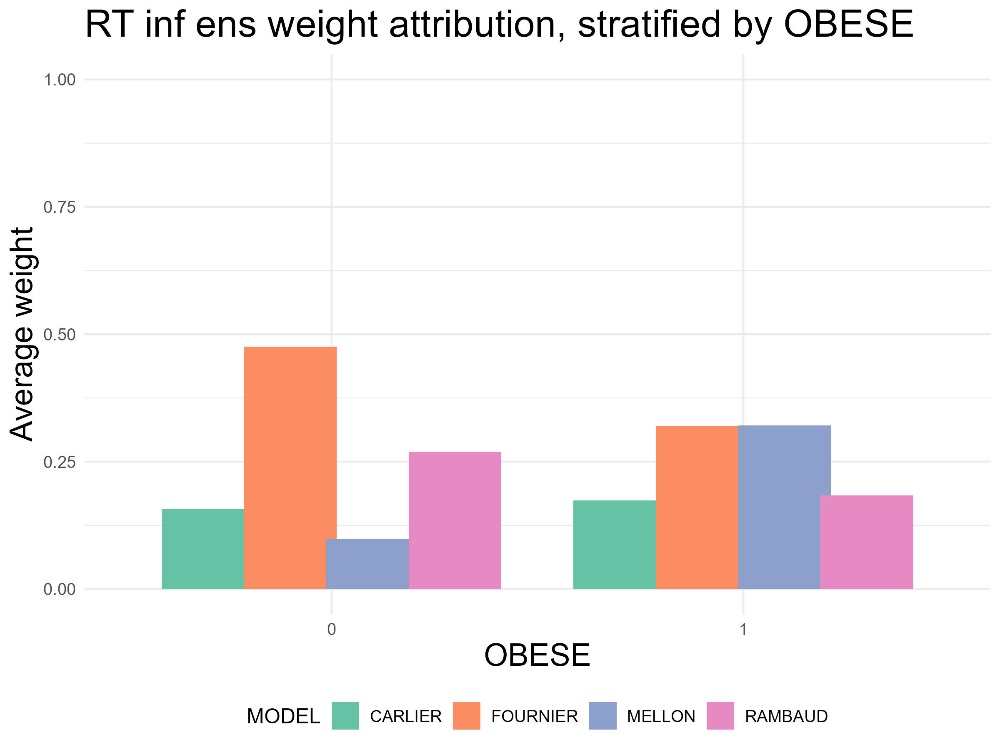

**Figure S10 a-b.** Average weight attribution to different models stratified by the OBESE categorical variable. FAMD (left) attributes weights solely based on similarity to the model cohorts without considering model performance, thus as as the Mellon cohort has the most obese subjects, for OBESE = 1, Mellon has the highest weight, and for the non-obese category (0) the weights are distributed between the three other models and Mellon has zero weight as it has zero non obese subjects. Although regression tree (RT) informed ensembling (on the right) takes into account model performance, it similarly attributed weight than FAMD showing the importance of model development cohort types over model performance.

#### FAMD

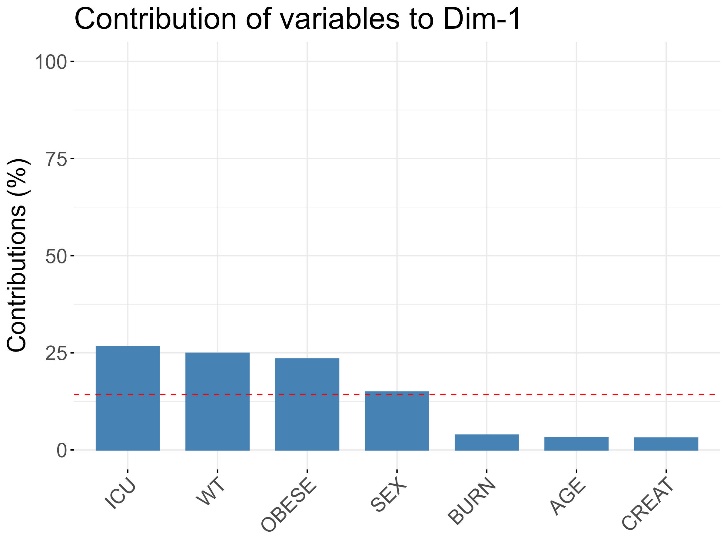

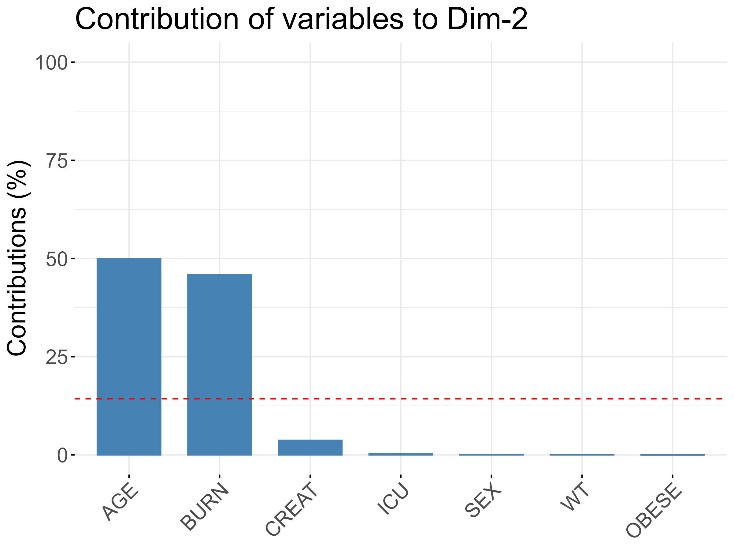

**Figure S11 a-b.** Contribution of covariates to the first (above) and second (below) principal component in FAMD. Notably, BURN contributed the least to the first dimension, and the most to the second.

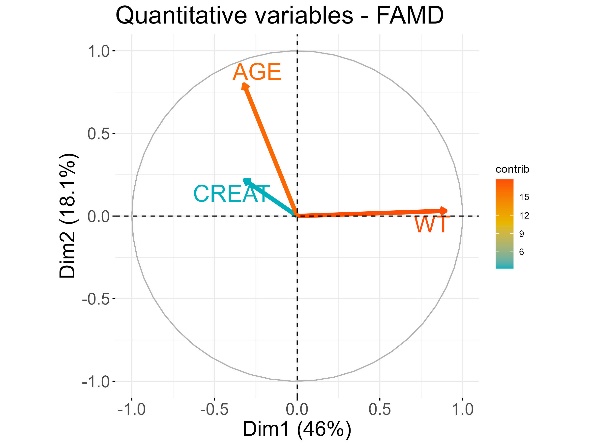

**Figure S12.** Contribution of continuous covariates to the first and second principal components in PCA. Body weight (WT) contributes the most to the results, and serum creatinine (CREAT) the least (of the continuous covariates). CREAT and age are positively correlated as the angle between them is smaller, while between CREAT and WT there is an important negative correlation.

#### WME

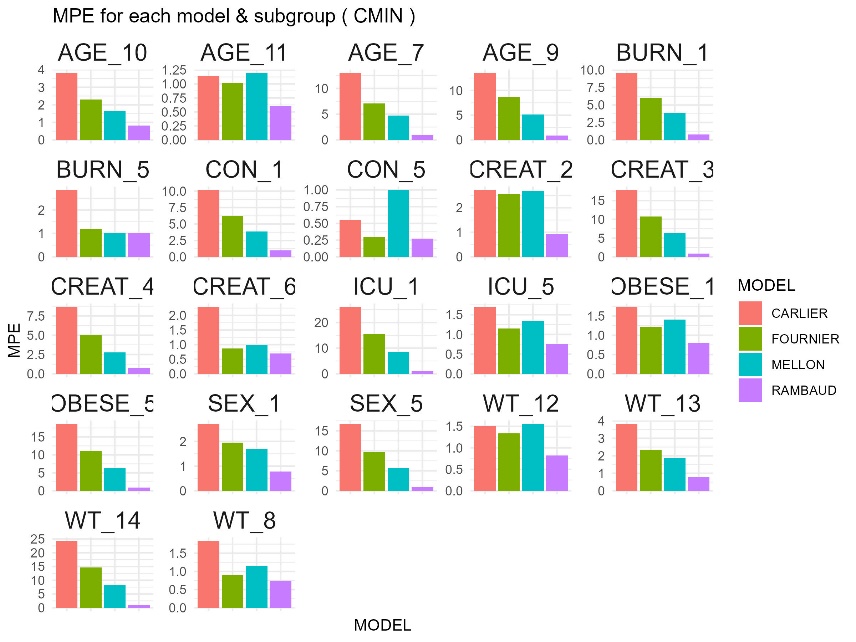

**Figure S13.** Mean percentage error (MPE) for each model in each covariate quantile/category.

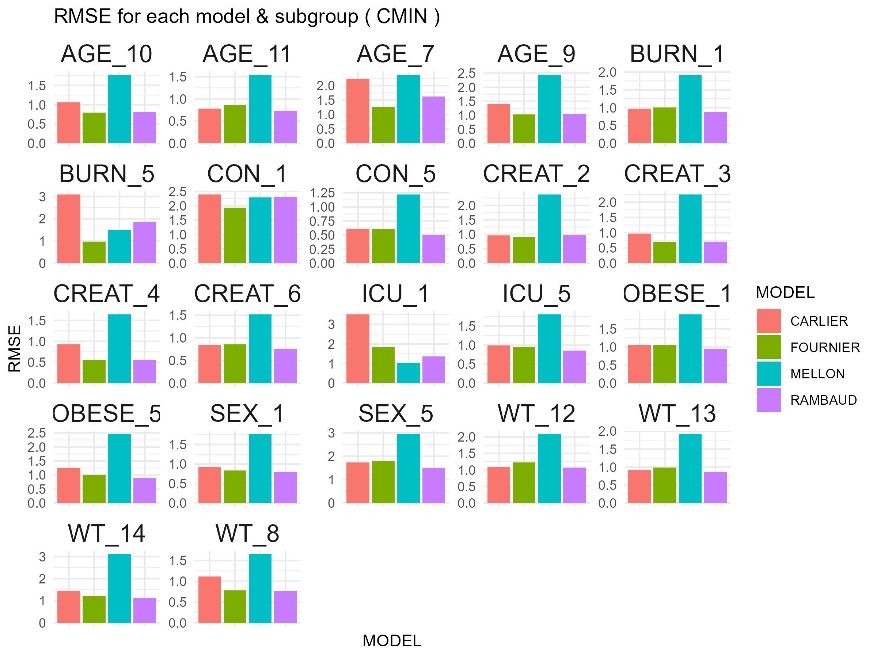

**Figure S14.** Relative root mean squared error (RMSE) for each model in each covariate quantile/category.

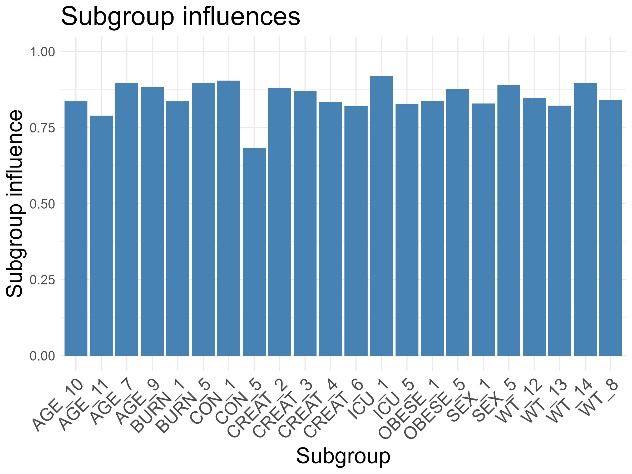

**Figure S15.** Subgroup influences (the proportion of incorrect predictions for each covariate quantile/category, all models included).

#### Machine Learning

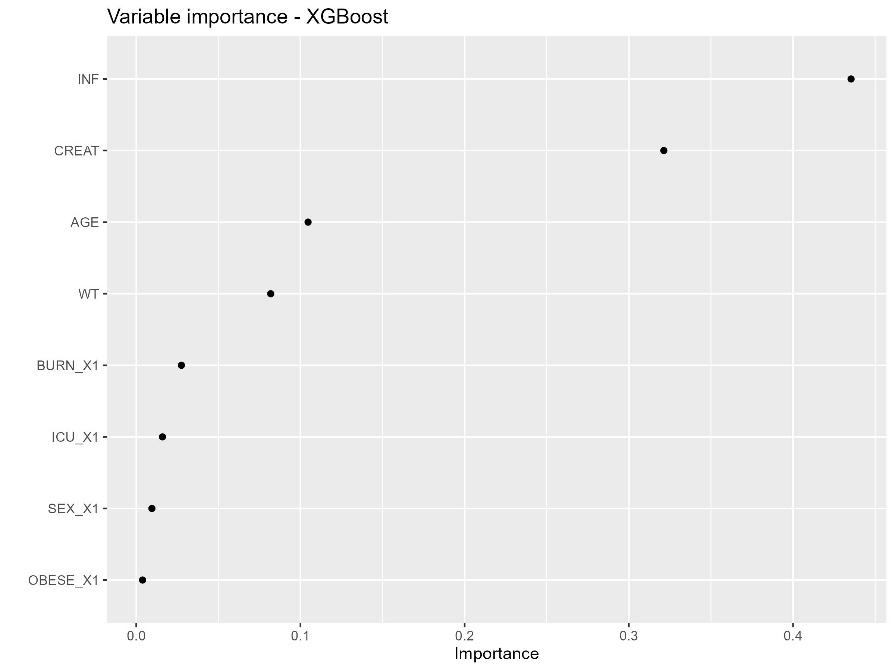

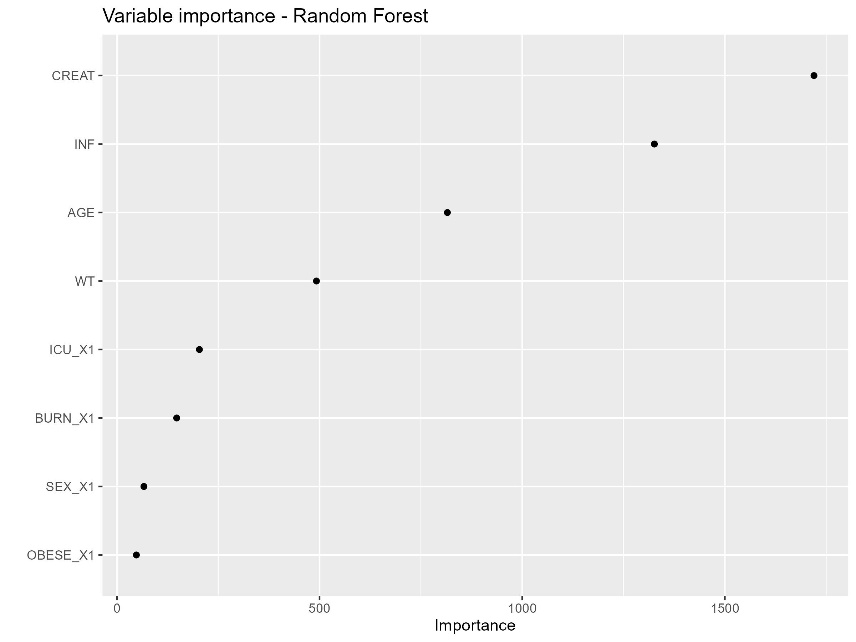

**Figure S16 a-b.** Impurity-based variable importance plot for XGBoost (left) and Random forest (right). For XGBoost, the most important predictor is the number of daily administrations (INF) and serum creatinine (CREAT) followed by being an Age and WT, while for Random forest, CREAT, INF and age contributed the most to the results.

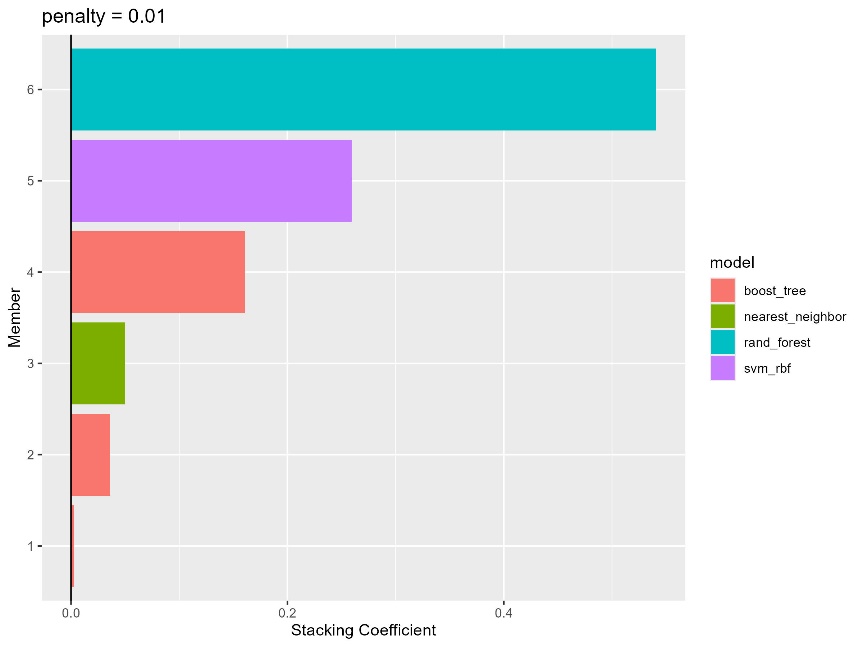

**Figure S17.** The contribution of different ML models to stacking. Random forest contributed the most to the model followed by support vector machine.

#### Method ensembling

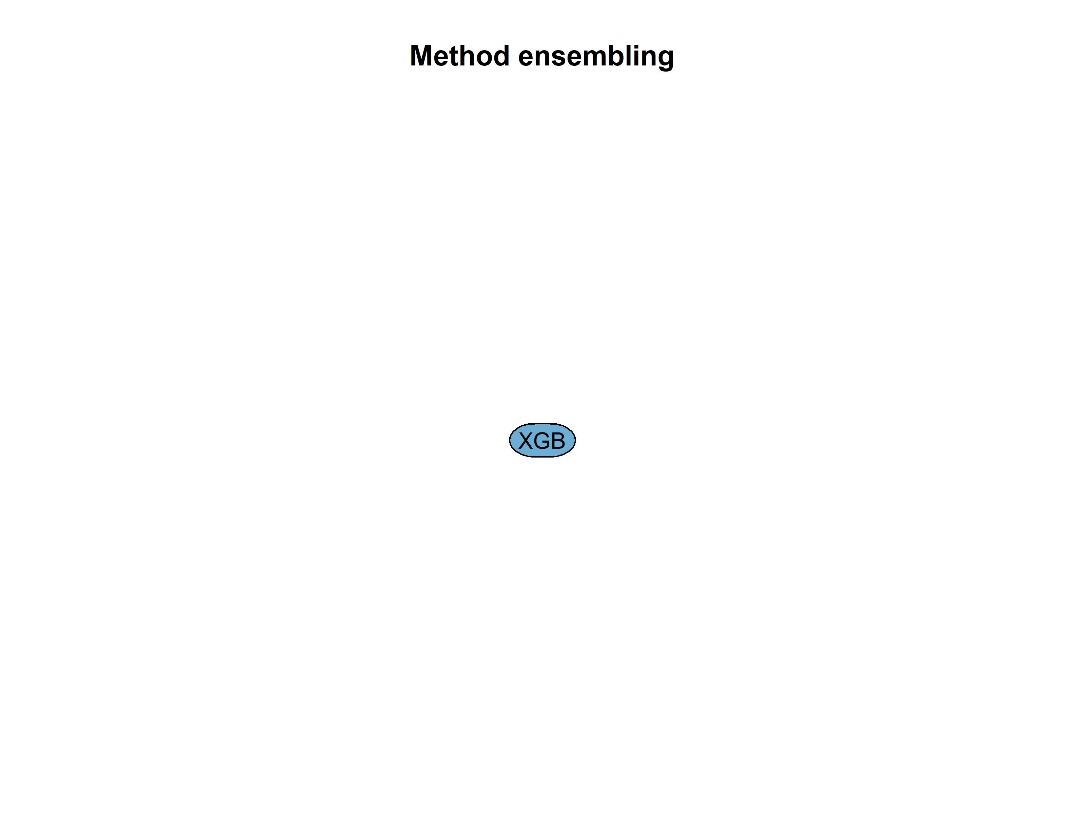

**Figure S18.** Decision tree for method ensembling. The predictors the covariates and the target is the best suited method for each subject. XGBoost was selected as the best-suited method for all subjects.

### Sensitivity analysis

**Table S3.** Sensitivity analysis for model inclusion on simulated data showing the percentage of correct dose predictions for each method.

| **Method** | **4 models (%)** | **Without Carlier (%)** | **Without Fournier (%)** | **Without Mellon (%)** | **Without Rambaud (%)** | **Without Mellon-Rambaud (%)** | **Without Fournier-Rambaud (%)** | **Without Fournier-Mellon (%)** | **Without Carlier-Rambaud (%)** | **Without Carlier-Mellon (%)** | **Without Carlier-Fournier (%)** |
| --- | --- | --- | --- | --- | --- | --- | --- | --- | --- | --- | --- |
| **Carlier** | 28 | | | | | | | | | | |
| **Fournier** | 32 | | | | | | | | | | |
| **Mellon** | 14 | | | | | | | | | | |
| **Rambaud** | 17 | | | | | | | | | | |
| **CT inf ensembling** | 31 | 34 | 34 | 38 | 21 | 29 | 25 | 45 | 21 | 49 | 38 |
| **RT inf ensembling** | 36 | 39 | 38 | 42 | 27 | 35 | 27 | 44 | 25 | 49 | 52 |
| **FAMD** | 37 | 45 | 42 | 40 | 28 | 32 | 27 | 45 | 27 | 49 | 48 |
| **XGBoost** | 36 | 37 | 39 | 37 | 27 | 26 | 28 | 43 | 28 | 44 | 47 |
| **SVM** | 39 | 42 | 41 | 39 | 30 | 28 | 28 | 46 | 31 | 45 | 51 |
| **RF** | 39 | 38 | 42 | 42 | 27 | 30 | 32 | 51 | 25 | 46 | 44 |
| **KNN** | 38 | 42 | 40 | 40 | 25 | 25 | 26 | 46 | 26 | 47 | 48 |

**Table S4.** Sensitivity analysis for model inclusion on clinical data showing the percentage of correct dose predictions for each method.

| **Method** | **4 models (%)** | **Without Carlier (%)** | **Without Fournier (%)** | **Without Mellon (%)** | **Without Rambaud (%)** | **Without Mellon-Rambaud (%)** | **Without Fournier-Rambaud (%)** | **Without Fournier-Mellon (%)** | **Without Carlier-Rambaud (%)** | **Without Carlier-Mellon (%)** | **Without Carlier-Fournier (%)** |
| --- | --- | --- | --- | --- | --- | --- | --- | --- | --- | --- | --- |
| **Carlier** | 40 | | | | | | | | | | |
| **Fournier** | 31 | | | | | | | | | | |
| **Mellon** | 20 | | | | | | | | | | |
| **Rambaud** | 28 | | | | | | | | | | |
| **CT inf ensembling** | 31 | 24 | 36 | 33 | 35 | 42 | 39 | 37 | 24 | 26 | 23 |
| **RT inf ensembling** | 36 | 27 | 37 | 40 | 40 | 40 | 39 | 37 | 22 | 28 | 28 |
| **FAMD** | 37 | 25 | 31 | 38 | 40 | 40 | 41 | 38 | 28 | 28 | 25 |
| **XGBoost** | 26 | 17 | 21 | 19 | 10 | 12 | 2 | 22 | 0 | 16 | 21 |
| **SVM** | 26 | 25 | 22 | 27 | 5 | 11 | 7 | 27 | 4 | 27 | 32 |
| **RF** | 25 | 23 | 27 | 26 | 5 | 5 | 5 | 27 | 0 | 28 | 19 |
| **KNN** | 26 | 27 | 27 | 24 | 3 | 2 | 7 | 27 | 1 | 26 | 29 |

**Table S5.** Sensitivity analysis for simulated dosing scheme repertoire showing the percentage of correct dose predictions for each method on simulated data.

| **Method** | **Based on model cohort (%)** | **Based on model cohort tested on intermittent** | **Based on model cohort tested on continuous** | **Intermittant 1 g Q8 (%)** | **Intermittant 1 g Q8 (%) tested on intermittent** | **Intermittant 1 g Q8 (%) tested on continuous** | **Continuous 10 g** | **Continuous 10 g tested on intermittent** | **Continuous 10 g tested on continuous** |
| --- | --- | --- | --- | --- | --- | --- | --- | --- | --- |
| **Carlier** | 28 |  |  | 28 |  |  | 28 |  |  |
| **Fournier** | 32 |  |  | 32 |  |  | 32 |  |  |
| **Mellon** | 14 |  |  | 14 |  |  | 14 |  |  |
| **Rambaud** | 17 |  |  | 17 |  |  | 17 |  |  |
| **CT inf ensembling** | 31 | 19 | 68 | 30 | 19 | 61 | 32 | 22 | 61 |
| **RT inf ensembling** | 36 | 25 | 68 | 32 | 23 | 58 | 31 | 23 | 53 |
| **WME** | 24 | 8 | 70 | 33 | 21 | 67 | 38 | 29 | 64 |
| **FAMD** | 37 | 27 | 68 | 37 | 27 | 68 | 37 | 27 | 68 |
| **XGBoost** | 36 | 28 | 60 | 19 | 25 | 0 | 23 | 8 | 69 |
| **SVM** | 39 | 28 | 70 | 17 | 22 | 0 | 21 | 6 | 67 |
| **RF** | 39 | 27 | 73 | 15 | 20 | 0 | 22 | 7 | 67 |
| **KNN** | 38 | 26 | 73 | 15 | 20 | 1 | 22 | 7 | 67 |

**Table S6.** Sensitivity analysis for simulated dosing scheme repertoire showing the percentage of correct dose predictions for each method on clinical data.

| **Method** | **Based on model cohort (%)** | **Based on model cohort tested on intermittent (%)** | **Based on model cohort tested on continuous (%)** | **Intermittant 1 g Q8 (%)** | **Intermittant 1 g Q8 tested on intermittent (%)** | **Intermittant 1 g Q8 tested on continuous (%)** | **Continuous 10 g (%)** | **Continuous 10 g tested on intermittent (%)** | **Continuous 10 g tested on continuous (%)** |
| --- | --- | --- | --- | --- | --- | --- | --- | --- | --- |
| **Carlier** | 40 |  |  | 40 |  |  | 40 |  |  |
| **Fournier** | 31 |  |  | 31 |  |  | 31 |  |  |
| **Mellon** | 21 |  |  | 21 |  |  | 21 |  |  |
| **Rambaud** | 28 |  |  | 28 |  |  | 28 |  |  |
| **CT inf ensembling** | 31 | 11 | 40 | 31 | 11 | 39 | 31 | 8 | 40 |
| **RT inf ensembling** | 36 | 14 | 45 | 30 | 0 | 43 | 32 | 3 | 45 |
| **WME** | 31 | 0 | 44 | 28 | 0 | 41 | 34 | 0 | 49 |
| **FAMD** | 35 | 3 | 49 | 35 | 3 | 49 | 35 | 3 | 49 |
| **XGBoost** | 26 | 5 | 35 | 7 | 8 | 6 | 34 | 30 | 36 |
| **SVM** | 26 | 11 | 32 | 0 | 0 | 0 | 38 | 35 | 39 |
| **RF** | 25 | 5 | 33 | 2 | 0 | 4 | 39 | 32 | 42 |
| **KNN** | 26 | 16 | 30 | 1 | 0 | 1 | 38 | 35 | 39 |

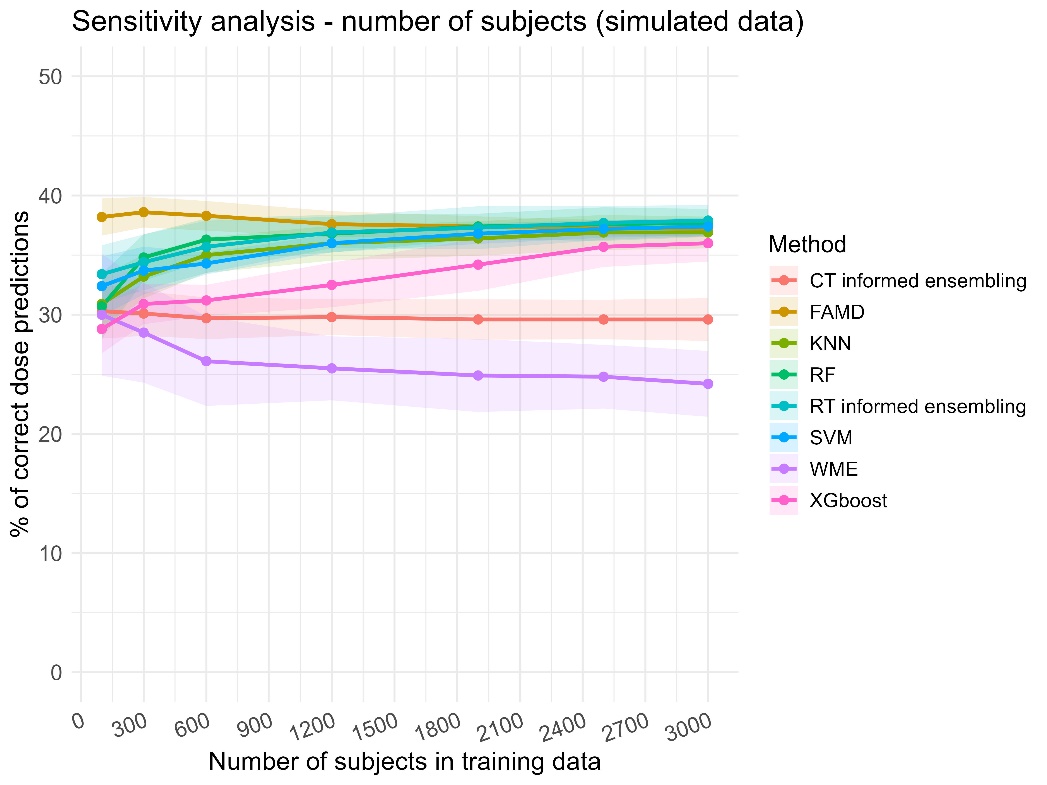

**Figure S19.** Sensitivity analysis for the number of simulated subjects in the training data on simulated data showing the percentage of correct dose predictions for each method and for different number of training subjects (from 100 to 3000) with shading indicating standard deviation calculated for 100 runs.

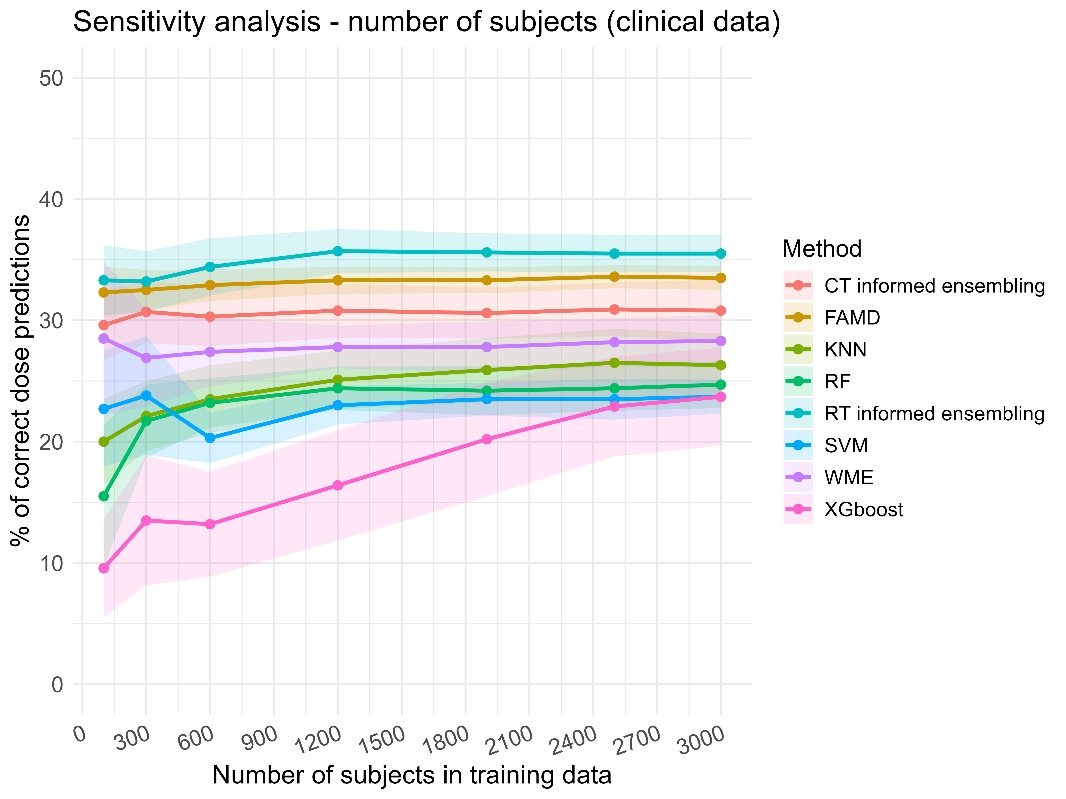

**Figure S20.** Sensitivity analysis for the number of simulated subjects in the training data on clinical data showing the percentage of correct dose predictions for each method and for different number of training subjects (from 100 to 3000) with shading indicating standard deviation calculated for 100 runs.

### **Bibliography**

1.	Johnson AEW, Bulgarelli L, Shen L, Gayles A, Shammout A, Horng S, et al. MIMIC-IV, a freely accessible electronic health record dataset. Scientific Data. 3 janv 2023;10(1):1.

2.	Silverman MP, Lipscombe TC. Exact Statistical Distribution of the Body Mass Index (BMI): Analysis and Experimental Confirmation. OJS. 2022;12(03):324‑56.

3.	Lepeule R, Bru JP, Canoui E, Gauzit R, Lesprit P, Jullien V, et al. Posologie standard et forte posologie : propositions du groupe de travail SPILF, SFPT & CA-SFM [Internet]. 2023 [cité 18 août 2025]. Disponible sur: https://www.infectiologie.com/UserFiles/File/spilf/recos/doses-spilf-sfpt-casfm-2023.pdf

4.	Rambaud A, Gaborit BJ, Deschanvres C, Le Turnier P, Lecomte R, Asseray-Madani N, et al. Development and validation of a dosing nomogram for amoxicillin in infective endocarditis. J Antimicrob Chemother. 1 oct 2020;75(10):2941‑50.

5.	Agema BC, Kocher T, Öztürk AB, Giraud EL, van Erp NP, de Winter BCM, et al. Selecting the Best Pharmacokinetic Models for a Priori Model-Informed Precision Dosing with Model Ensembling. Clin Pharmacokinet. oct 2024;63(10):1449‑61.

6.	Pagès J. Analyse factorielle de données mixtes. Revue de Statistique Appliquée. 2004;52(4):93‑111.

7.	Reprint of: Mahalanobis, P.C. (1936) « On the Generalised Distance in Statistics. » Sankhya A. déc 2018;80(S1):1‑7.

8.	Kuhn M, Wickham W. Tidymodels: a collection of packages for modeling and	machine learning using tidyverse principles. [Internet]. 2020. Disponible sur: https://www.tidymodels.org
